# Stroke induces early recurrent vascular events by inflammasome-dependent atherosclerotic plaque rupture

**DOI:** 10.1101/2023.02.01.526550

**Authors:** Jiayu Cao, Stefan Roth, Sijia Zhang, Anna Kopczak, Marios K. Georgakis, Xinghai Li, Alexander Dutsch, Thomas G Liman, Matthias Endres, David Brough, Jack P. Green, Saskia Wernsdorf, Christina Fürle, Olga Carofiglio, Jie Zhu, Yaw Asare, DEMDAS Study Group, Martin Dichgans, Hendrik B. Sager, Gerrit M. Große, Arthur Liesz

## Abstract

The risk of early recurrent events after stroke remains high despite currently established secondary prevention strategies. Risk is particularly high in patients with atherosclerosis, with more than 10% of patients experiencing early recurrent events. However, despite the enormous medical burden of this clinical phenomenon, the underlying mechanisms leading to increased vascular risk and recurrent stroke are largely unknown. Here, using a novel mouse model of stroke-induced recurrent ischemia, we show that stroke leads to activation of the AIM2 inflammasome in vulnerable atherosclerotic plaques via an increase of circulating cell-free DNA from the ischemic tissue. Enhanced plaque inflammation post-stroke results in plaque destabilization and atherothrombosis, finally leading to arterio-arterial embolism and recurrent stroke within days after the index stroke. We confirm key steps of plaque destabilization also after experimental myocardial infarction and in carotid artery plaque samples from patients with acute stroke. Neutralization of cell-free DNA by DNase treatment or inhibition of inflammasome activation reduced the rate of stroke recurrence after experimental stroke. Our findings present an explanation for the high recurrence rate after incident ischemic events in atherosclerotic patients. The detailed mechanisms uncovered here provide so far clinically uncharted therapeutic targets for which we show high efficacy to prevent recurrent events. Targeting DNA-mediated inflammasome activation after remote tissue injury represents a promising avenue for further clinical development in the prevention of early recurrent events.

## Introduction

Stroke is the second leading cause of death worldwide^1^ and a leading cause of long-term disability with incidence rates rising due to demographic aging^2^. An important factor contributing to this sociomedical burden is the high risk of recurrent vascular events such as stroke and myocardial infarction. A systematic review and meta-analysis reported a pooled recurrent stroke risk of 11.1% at 1 year^3,4^. Importantly, the risk of myocardial infarction is likewise substantially increased in stroke survivors^5^. The precise mechanisms underlying this increased vascular risk after an ischemic event are unclear. It is equally unexplained why the hazards for recurrence are time-dependent and particularly pronounced in the early phase after the event. Previous epidemiological studies have indicated that vascular risk is particularly increased in patients with large-artery atherosclerosis and to a lesser degree in patients with other stroke etiologies^6^. This suggests an abundant residual vascular risk in patients with atherosclerosis, which is not effectively prevented by current secondary prevention measures.

Atherosclerosis is a chronic inflammatory disease, where immune cells in atherosclerotic plaques contribute to the progression and evolving vulnerability^7^. We and others have previously shown that stroke leads to a systemic inflammatory response, which can contribute to progression of vascular inflammation and plaque load in established atherosclerosis^8–10^. Epidemiological data on inflammatory biomarkers and atheroprogression in stroke patients likewise suggest that inflammation can be associated with progression of atherosclerosis and even with recurrent ischemic events^11,12^.

Based on these findings, we formulated the hypothesis that stroke might facilitate the occurrence of subsequent vascular events by inflammatory mechanisms. However, the detailed mechanisms along this supposed brain-immune-vascular axis are currently unknown and therefore specific immunological targets to potentially reduce the rate of early recurrence after stroke are missing. A potential reason for this lack of information on such a pressing biomedical question might be the lack of suitable animal models to study recurrent ischemic events. Commonly used models for atherosclerosis in rodents differ from the situation in high-risk patients with cardiovascular disease in that the atherosclerotic plaques are less complex, not rupture-prone and barely affect the cerebrovascular circulation.

Here, we developed an adapted mouse model of rupture-prone high-risk plaques of the carotid artery in combination with contralateral experimental stroke or with myocardial infarction (MI). We found that both stroke and MI induced a destabilization of atherosclerotic plaques, leading to recurrent ischemic events. We identified DNA-mediated activation of the AIM2 inflammasome as the immunological mechanism leading to plaque destabilization via matrix metalloproteinase activation, finally leading to atherothrombosis and arterial embolism. The same pathophysiological processes were confirmed in human atherosclerotic plaques obtained from patients within the first days after an acute ischemic stroke. Targeting this immunological pathway efficiently prevented recurrent ischemic events after experimental stroke.

## Results

### Incident stroke induces early recurrent events in atherosclerosis

Since previous epidemiological data on stroke recurrence by stroke etiology are more than 20 years old^6^ and clinical practice has substantially changed in this time period, we assessed the recurrence rate of cerebrovascular events (stroke and transient ischemic attack) using pooled data (n=1798 stroke patients) from two ongoing clinical cohorts with information on etiology of the first stroke according to TOAST criteria^13,14^. Focusing on the early phase after stroke (days 0-30 after the index event) recurrences rates were markedly higher in patients with large artery atherosclerotic (LAA) stroke compared to other stroke etiologies and nearly as high as for the entire period from days 31-360 combined (**Fig. 1A**, Suppl. Fig. 1A). This indicates that current strategies for secondary stroke prevention fail to efficiently reduce early recurrent events in patients with LAA stroke. This clinical notion was further confirmed by treating atherosclerotic mice undergoing experimental stroke with high dose rosuvastatin and aspirin—the current standard scheme for secondary stroke prevention in patients. This high-dose treatment had no effect on mortality, vascular inflammation and plaque load within the first week after experimental stroke (Suppl. Fig. 1B-F), confirming the inefficiency of current secondary prevention therapy on early atherosclerotic plaque progression within the first week after stroke. To study the mechanisms underlying the increased risk of recurrent events in atherosclerotic patients, we established an animal model of unilateral highly stenotic and hemodynamically relevant atherosclerotic plaques of the common carotid artery (CCA). This involved the induction of turbulent flow by a stenotic tandem ligation (Suppl. Fig. 2). We used this model to test the effects of experimentally induced acute ischemic stroke in the hemisphere contralateral to the CCA tandem stenosis (TS) on atherosclerotic plaque morphology and destabilization (**Fig. 1B**). We screened for the occurrence of secondary ischemic events in the brain hemisphere supplied by the stenotic atherosclerotic CCA (i.e. contralateral to the experimentally induced stroke) by magnetic resonance imaging (MRI) and histological analysis of cell loss (Fluoro Jade C and TUNEL staining) and microgliosis (**Fig. 1C**). We found that experimental stroke resulted in secondary brain ischemia in 30% of the animals, while this was not observed in any of the animals with CCA TS and sham surgery (**Fig. 1C**). This observation suggested stroke-induced rupture of remote CCA plaques leading to secondary stroke events. This hypothesis was further supported by the presence of intravascular thrombi in the supplying arteries (e.g. the anterior cerebral artery) proximal to the secondary stroke territory (in **Fig. 1D**). Again, we found no such thrombi in sham-operated animals. Correspondingly, the plaque vulnerability index for the stenotic CCA (a previously established score of several morphological markers indicating plaque risk to rupture^15^, Suppl. Fig. 3A-D) was significantly increased in mice after experimental stroke and was even further exacerbated in mice with secondary events after the primary stroke (**Fig. 1E**). Histological indication of plaque rupture was observed in 40% of mice after stroke compared to 10% of mice undergoing sham surgery (Suppl. Fig. 3E). We further validated the concept of secondary remote plaque rupture in a model of myocardial infarction (MI). Similar to stroke, MI increased vulnerability of the remote CCA plaque (**Fig. 1F**, Suppl. Fig. 4). Flow cytometry analysis of CCA plaques revealed increased cellular plaque inflammation after stroke in comparison to sham surgery (Suppl. Fig. 5). We further analyzed the mechanism of enhanced immune cellularity either by invasion or proliferation using antibody labeling of circulating immune cells or histological assessment of the proliferation marker Ki67, respectively. We found that both the *de novo* recruitment of circulating leukocytes into plaques and local proliferation of leukocytes were increased after stroke (**Fig. 1G-I**) and that the local proliferation rate was associated with the occurrence of recurrent events (**Fig. 1I**).

**Figure 1.**
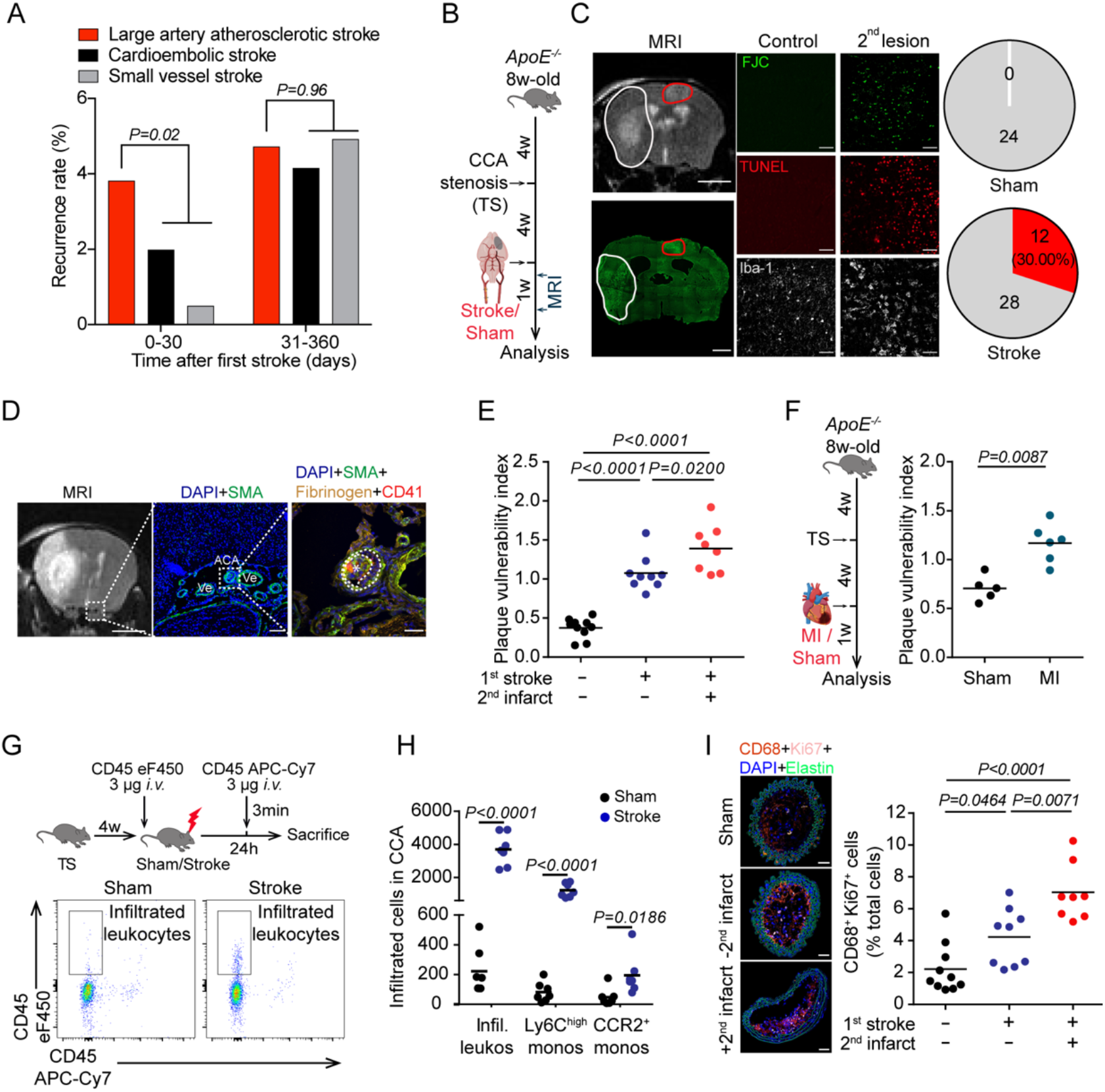
Ischemic events induce recurrent stroke and exacerbate plaque vulnerability. (**A**) Recurrence rates by incident stroke etiology in the first month (days 0-30) and months 2-12 (days 31-360) in a population of 1798 stroke patients in Germany (p values derived from log-rank test in Kaplan-Meier curves, Suppl. Fig. 1A). (**B**) Experimental design: 8-week-old ApoE^−/-^ mice fed a high cholesterol diet (HCD-fed *ApoE^−/−^*) underwent tandem stenosis (TS) surgery, and stroke or Sham surgery 4 weeks later. The recurrence of secondary ischemia in the contralateral hemisphere was examined by MRI (days 2 and 7) and histological analysis 7d after sham or stroke surgery. (**C**) Representative MRI image with a recurrent stroke (white dashed line: primary stroke area; red dotted line: recurrent contralateral stroke). The lower left panel shows Fluoro Jade C (FJC) staining corresponding to the MRI (scale bars =3 mm). Right images: representative images of histological stainings (FJC, TUNEL, Iba-1) from control mice or mice with a secondary lesion (scale bar=50 μm). Pie charts illustrate stroke recurrence rate 7d after sham or stroke surgery (sham: n=24, stroke: n=40; red, with secondary lesion; grey, without secondary lesion). (**D**) Representative images of a thrombus found in the anterior cerebral artery (ACA). Ve = venules. * indicates the thrombus within the ACA. (**E**) Quantification of plaque vulnerability in CCA sections 7d after sham or stroke surgery (ANOVA, n= 8-10 per group). (**F**) Experimental design as shown in (B), but with induction of MI instead of Stroke, and quantification of plaque vulnerability (U test, n=5-6 per group). (**G**) Experimental design and representative FACS plots for (**H**) quantification of total infiltrated leukocytes and Ly6C^high^ and CCR2^+^ monocyte subpopulations in CCA 24h after sham or stroke surgery (U test, n=7 per group). (**I**) Representative images (left) and quantification (right) of proliferating macrophages between groups as in (E) (ANOVA, n=8-10 per group).

### Post-stroke DNA release induces AIM2 inflammasome activation in atherosclerotic plaques

Further analyses of the inflammatory milieu in atherosclerotic CCA plaques revealed a substantial increase in local IL-1β production after stroke, suggesting a stroke-induced inflammasome activation within atherosclerotic plaques (**Fig. 2A**). Local inflammasome activation was confirmed by Western blot analysis of caspase-1 cleavage–the effector enzyme of the inflammasome–in the atherosclerotic plaque and independently verified by histological analysis with increased caspase-1 expression in CCA plaques after stroke (Suppl. Fig. 6). Blocking inflammasome activity by a specific caspase-1 inhibitor (VX765) prevented the proliferation of immune cells within CCA plaques after stroke and attenuated the invasion of pro-inflammatory circulating monocytes (**Fig. 2B, C**). Consequently, inflammasome inhibition significantly reduced plaque vulnerability after experimental stroke to levels comparable to sham-operated mice (**Fig. 2D** and Suppl. Fig. 7). We next investigated which specific inflammasome mediates this effect of stroke on CCA plaque exacerbation by applying specific pharmacological inhibitors of the NLRP3 and of the AIM2 inflammasome^16^. We focused on these two abundant inflammasome subtypes, because the NLRP3 inflammasome has previously been implicated in the development of atherosclerosis, while stroke leads to systemic AIM2 inflammasome activation^17,18^. Interestingly, only the inhibition of the AIM2 but not of the NLRP3 inflammasome prevented inflammasome activation in the CCA plaque after stroke (**Fig. 2E, F** and Suppl. Fig. 8A). Moreover, we observed a transient increase in the serum concentration of cell-free DNA in the acute phase after stroke and MI, which is the ligand of the AIM2 inflammasome, suggesting that release of cell-free DNA after stroke and MI could be the mediator linking remote organ ischemia to the exacerbation of plaque inflammation (**Fig. 2G** and Suppl. Fig. 8B, C). In order to test this hypothesis, we injected DNA into mice with a CCA stenosis without stroke or sham surgery and observed increased plaque inflammasome activation similar to the condition after stroke (**Fig. 2H, I** and Suppl. Fig. 8D, E). In contrast, neutralization of cell-free DNA after stroke by therapeutic administration of recombinant DNase following stroke induction again significantly reduced the plaque inflammasome activation, as measured by caspase-1 cleavage and IL-1β secretion (**Fig. 2J, K**). In summary, we identified the activation of the AIM2 inflammasome in vulnerable atherosclerotic plaques by stroke-induced DNA release and subsequent plaque rupture as the cause of recurrent ischemic events.

**Figure 2.**
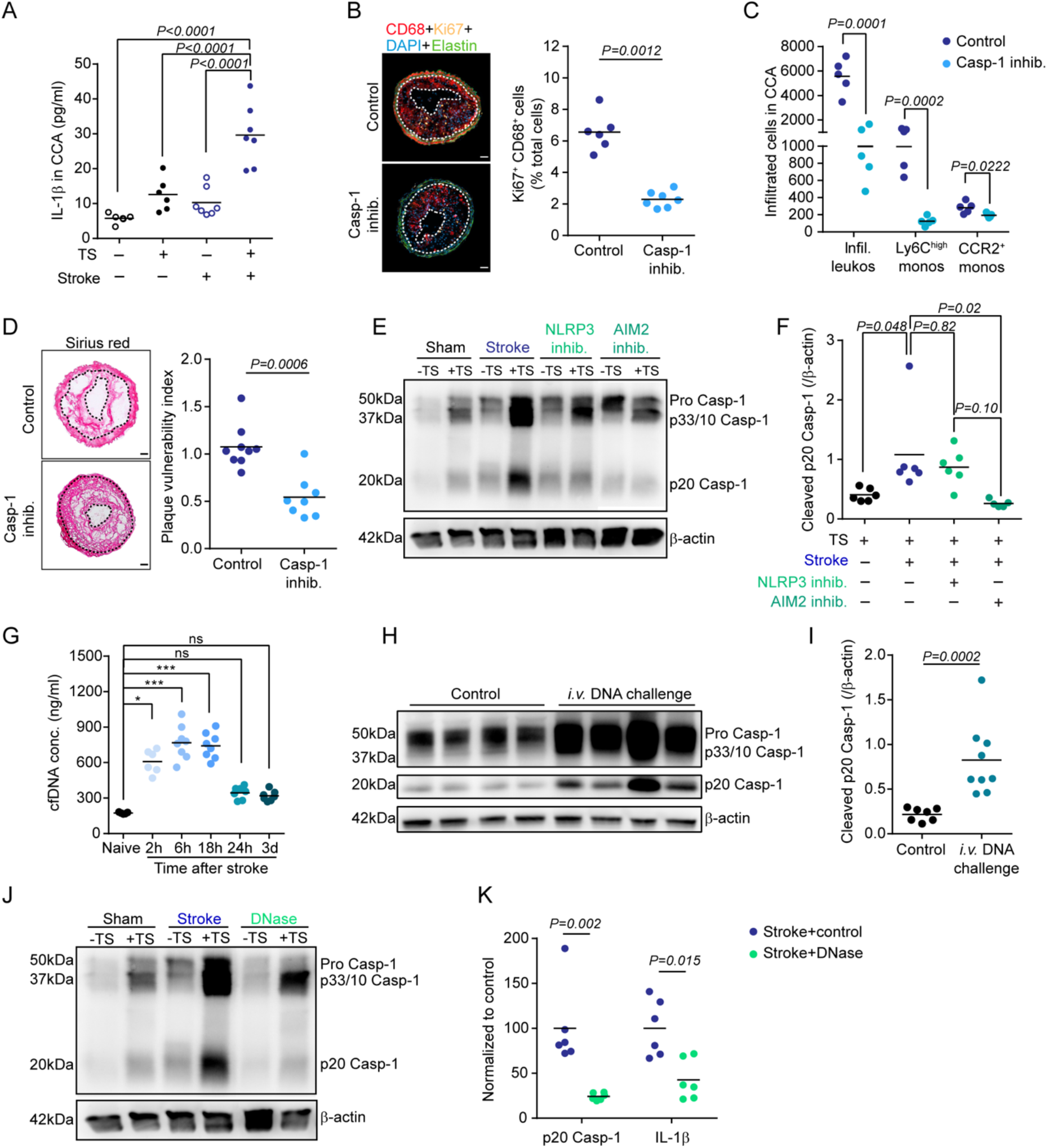
Stroke induces dsDNA-dependent inflammasome activation in atherosclerotic plaques. (**A**) ELISA analysis of IL-1β concentrations in murine CCA lysates 24h after sham or stroke surgery (ANOVA, n=6-7 per group). (**B**) Representative images (left) and quantification of proliferating macrophages normalized to total cell counts (right; U test, n=6-7 per group) in CCA sections of control- or caspase-1 inhibitor (VX765)-treated mice 7d after stroke. (**C**) Flow cytometry analysis of infiltrated leukocyte counts between treatment groups (corresponding to experiment in 1G). (**D**) Representative picro sirius red stainings of CCA from mice which received control or caspase-1 inhibitor treatment 1 w after stroke (left; scale bar = 50 μm). Right: Corresponding quantification of plaque vulnerability (U test, n=8-9 per group). (**E**) Representative immunoblot micrograph of the different cleavage forms of caspase-1 (Casp-1) in CCA lysates 24h after sham, stroke, stroke + NLRP3 inhibitor (MCC950) or AIM2 inhibitor (4-sulfonic calix[6]arene) administration. (**F**) Corresponding immunoblot quantification of cleaved p20 Casp-1 intensity normalized to β-actin (ANOVA test, n=5-6 per group). (**G**) Total cell-free DNA (cfDNA) serum concentrations at the indicated time points after experimental stroke in comparison to naïve mouse serum (U test, n=5-8 per group). (**H**) Representative immunoblot micrograph and (**I**) quantification of cleaved p20 Casp-1 intensity normalized to β-actin in CCA lysates from HCD-fed *ApoE^−/−^* with TS surgery or 24h after *i.v*. DNA challenge (U test, n=7-9 per group). (**J, K**) HCD-fed *ApoE^−/−^* mice with TS surgery received 1000U i.v. DNase immediately before stroke. Cleaved p20 Casp-1 and IL-1ß were quantified in CCA lysates 24h later by immunoblot (U test, n= 6). Data is shown as a ratio of the DNase treated group to the mean of the control group.

### Inflammatory plaque degradation by matrix metalloproteinases leads to atherothrombosis

Conceivably, enhanced plaque inflammation—as observed here after stroke induction—could contribute to plaque rupture and secondary arterio-arterial embolism resulting in secondary infarctions. However, the exact mechanisms linking plaque inflammation to plaque destabilization and atherothrombosis are poorly studied. We found that stroke resulted in an increased activity of matrix metalloproteinases (MMPs) as detected by *in situ* zymography of CCA plaque sections and validated by gel zymography for MMP2 and MMP9 (**Fig. 3A** and Suppl. Fig. 9A, B). To test whether soluble blood mediators mediate this effect on MMP activity after stroke, we treated bone-marrow derived macrophages (BMDM) with serum obtained from mice 4h after stroke or sham surgery. Conditioning of BMDMs with serum from stroke mice resulted in massively increased active MMP2 and MMP9 secretion in comparison to sham serum treatment, both in total protein content (western blot) and enzymatic activity (zymography), suggesting a causative role of circulating factors after stroke on MMP expression and activation in the CCA plaque (**Fig. 3B**, quantification in Suppl. Fig. 9C-E). Given our finding of a critical role of AIM2 inflammasome activation, we tested the influence of IL-1β derived from inflammasome activation on MMP activity by treating BMDMs with increasing doses of recombinant IL-1β. Indeed, we detected a dose-dependent effect of IL-1β on MMP expression (**Fig. 3C** and Suppl. Fig. 9F). While MMP-mediated degradation of the extracellular matrix causes plaque destabilization^19^, formation of the actual thrombus on ruptured plaques depends on activation of the coagulation cascade, particularly by the contact activation pathway initiated by Factor XII (F.XII) exposition to damaged tissue surfaces^20^. We therefore performed *en face* staining of the whole CCA and compared activated Factor XII (F.XIIa) deposition on the luminal surface at the area of the CCA plaque between stroke or sham operated mice. We found a significant increase in F.XIIa deposition on the plaque surface after stroke (**Fig. 3D**), which was confirmed by western blot analysis of whole CCA plaques (**Fig. 3E** and Suppl. Fig. 10A). Interestingly, an intravenous DNA challenge without stroke mimicked the effect of stroke with increased F.XIIa deposition at the CCA plaque (**Fig. 3F** and Suppl. Fig. 10B). These findings suggest that the AIM2 inflammasome activation and resulting IL-1β release after stroke could lead to plaque destabilization and atherothrombosis via activation of plaque-degrading MMPs and subsequent F.XIIa deposition, which in turn could trigger thrombus formation. Correspondingly, *in vivo* inflammasome inhibition using the caspase-1 inhibitor VX765 significantly reduced plaque MMP activity and luminal F.XIIa deposition (**Fig. 3G, H** and Suppl. Fig. 10C). Similarly, i.v. DNase treatment significantly reduced MMP activity and F.XIIa deposition (**Fig. 3I** and Suppl. Fig. 10D).

**Figure 3.**
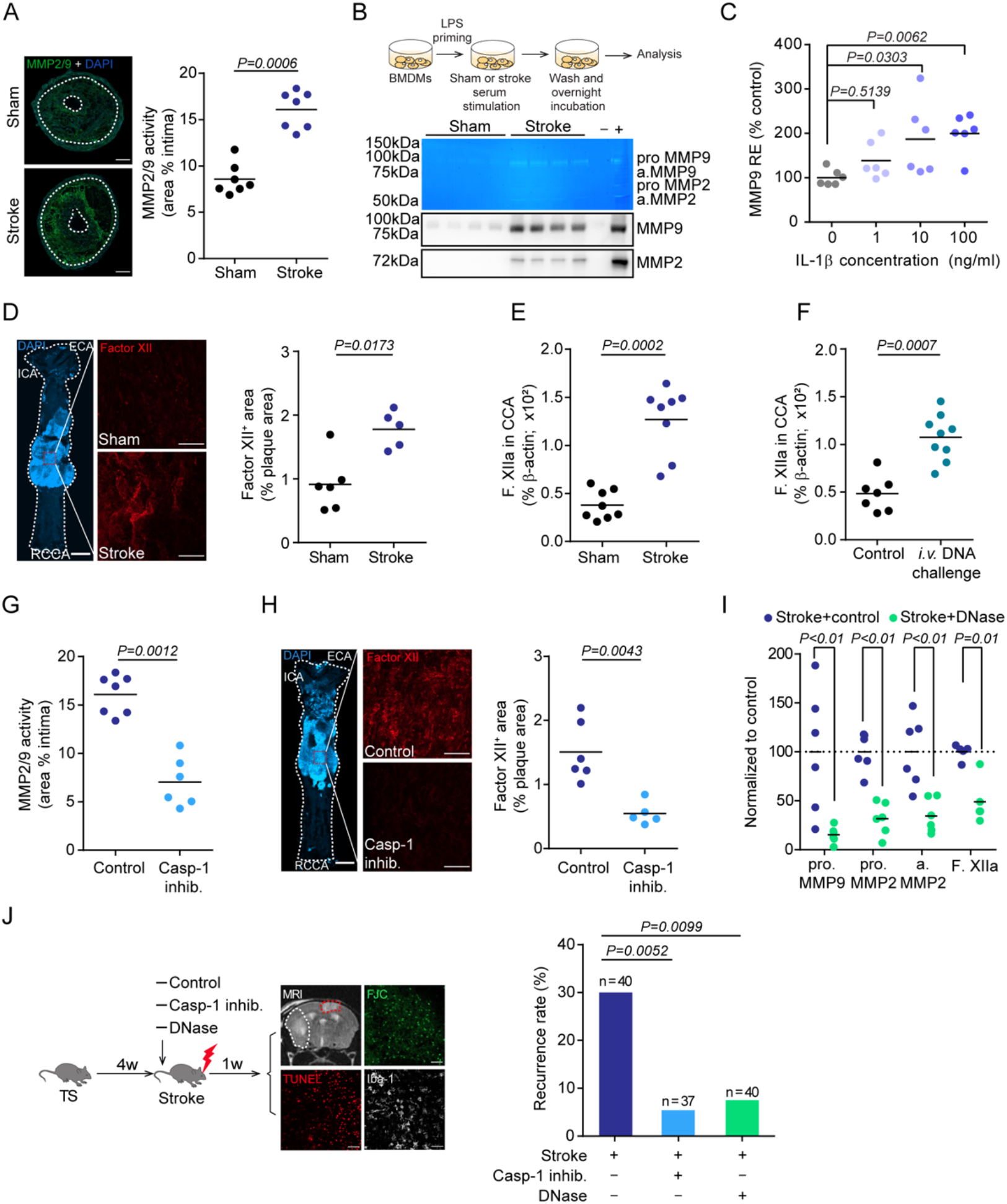
Inhibition of post-stroke inflammasome activation prevents plaque destabilization and recurrent stroke events. (**A**) Representative images quantification of MMP2/9 activity by *in situ* zymography on CCA sections 7d after sham or stroke surgery (U test, n= 7 per group). (**B**) Schematic experimental design and representative images for gelatin zymography (top) and immunoblot micrograph (bottom) of MMP9 and MMP2 in the culture medium (quantification in Suppl. Fig. 8). (**C**) Relative expression (RE) for MMP9 mRNA in BMDMs after IL-1ß stimulation (H test, n=6 per group). (**D**) Whole CCA *en face* immunofluorescence staining of F.XII and quantification 24h after stroke or sham surgery (U test, n=5-6 per group). (**E**) Quantification of activated F.XII (F.XIIa) in CCA lysates 7d after sham or stroke surgery (U test, n=8 per group). (**F**) Quantification of F.XIIa in CCA lysates 24h after *i.v*. DNA challenge (U test, n=7- 9 per group). (**G**) Quantification of *in situ* zymography for MMP2/9 activity in CCA sections 7d after stroke (U test, n=6-7 per group). (**H**) Quantification of F.XII+ area on CCA *en face* images, corresponding to (D), between treatment groups (U test, n=5-6 per group). (**I**) ProMMP9, proMMP2, activated MMP2 (a.MMP2) and F.XIIa were quantified in mice receiving 1000U DNase after stroke and normalized to control-treated mice (U test, n= 5-6 per group). (**J**) Left: Schematic illustration of experimental design for therapeutic intervention. Right: Quantification of 7d recurrence rate between groups by the same analysis strategy as in Fig. 1B,C (chi-square test).

### Blocking DNA-mediated inflammasome activation prevents early stroke recurrence

Finally, we analyzed whether blocking DNA-mediated inflammasome activation after stroke could be used therapeutically to prevent recurrent ischemic events. For this, mice with CCA tandem stenosis were treated with the caspase-1 inhibitor VX765 or recombinant DNase. The occurrence of secondary ischemic lesions was analyzed 7d after induction of the primary stroke and in the hemisphere contralateral to the primary stroke i.e. within the territory supplied by the stenotic CCA. Indeed, both caspase-1 inhibition and DNase treatment greatly and significantly reduced the recurrence rate (**Fig. 3J**). Notably, this therapeutic effect was achieved in a large sample size of animals (total sample size: 117 mice) and translates to a relative risk reduction of 82% and 75% for VX765 and DNase, respectively.

To further validate the translational relevance of the identified mechanisms, we obtained carotid endarterectomy samples from highly stenotic carotid artery plaques of asymptomatic patients and patients during the acute phase of ischemic stroke (**Fig. 4A**). Flow cytometry analysis of the plaque material revealed a significant increase in monocyte counts in plaques from symptomatic compared to asymptomatic patients (**Fig. 4B**), while blood monocyte and lymphocyte counts did not differ (Suppl. Fig. 11). Correspondingly, we found a significant increase in circulating cell-free DNA in plasma of stroke patients, as well as a substantial increase in markers of inflammasome priming (pro-caspase-1 expression) and inflammasome activation (cleaved p20 isoform of caspase-1) in plaque material (**Fig. 4C-F**). We observed a similar increase in cell-free DNA in patients after myocardial infarction, emphasizing again the generalizability of our findings across ischemic organ damage (**Fig. 4G**). Notably, we also detected a more than 10-fold increase in MMP9 activity by gel zymography of plaques from symptomatic compared to asymptomatic patients (**Fig. 4H**, Suppl. Fig. 11). Finally, the amount of plaque-associated F.XIIa was significantly increased in atherosclerotic plaques after stroke (**Fig. 4I**). Hence, the analyses of human endarterectomy samples confirmed all key steps of DNA-mediated inflammasome activation, vascular inflammation, MMP activation and initiation of thrombus formation in stroke patients as identified in our animal model.

**Figure 4.**
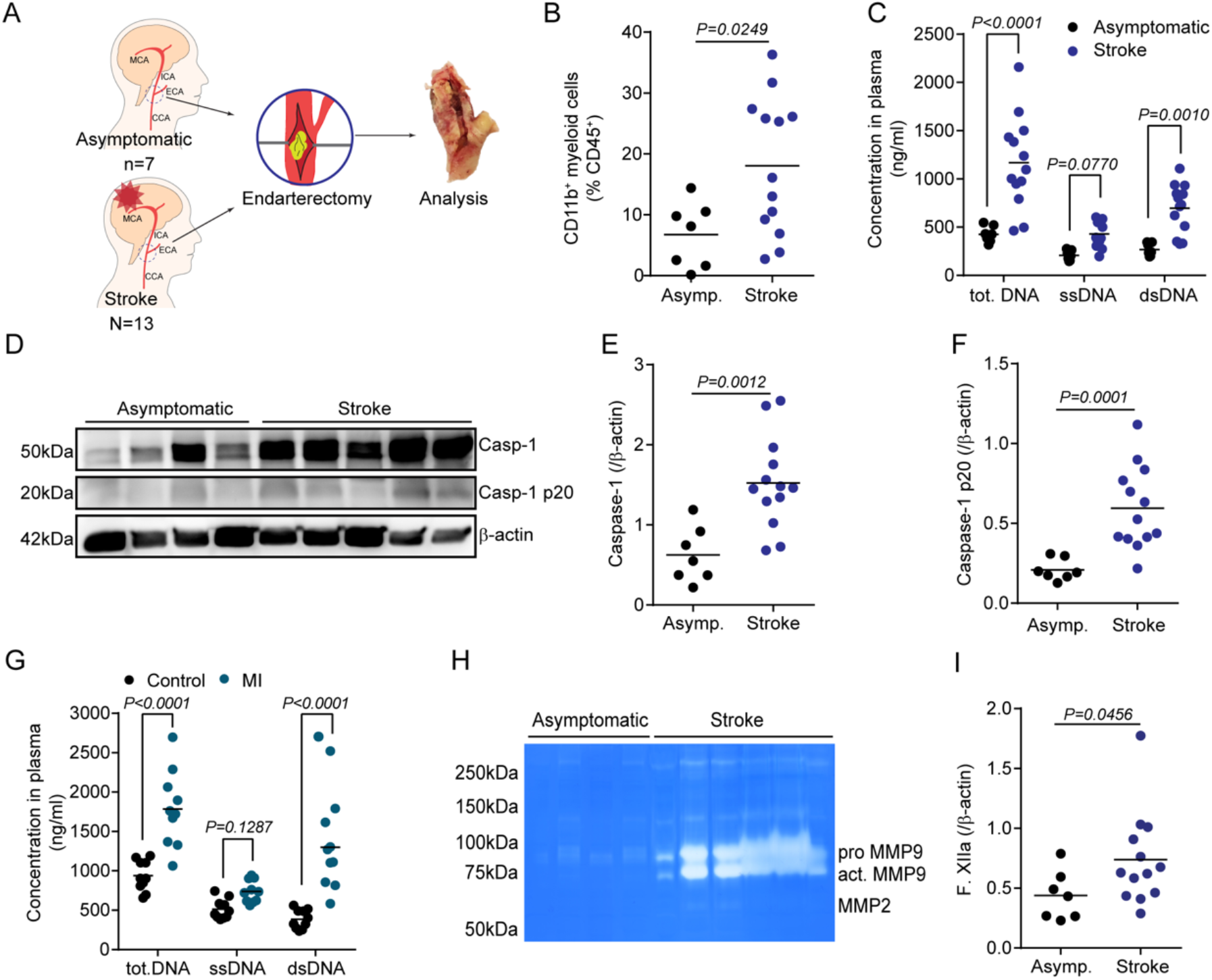
Stroke increases atherosclerotic plaque inflammation and MMP activity in patients. (**A**) Study layout illustrating collection of endarterectomy samples from 7 asymptomatic patients with internal carotid artery stenosis and from 13 stroke patients undergoing endarterectomy of the symptomatic carotid artery in the acute phase after stroke. (**B**) Flow cytometry analysis of plaques showing the percentage of CD11b^+^ myeloid cells out of total leukocytes (CD45^+^). (**C**) Quantification of total cell-free DNA (tot.DNA), single-stranded DNA (ssDNA) and double-stranded DNA (dsDNA) in plasma. (**D**) Representative immunoblot micrograph of the different cleavage forms of caspase-1 (Casp-1) in atherosclerotic plaque lysates from asymptomatic or symptomatic patients and (**E, F**) the corresponding quantification of total Casp-1 and cleaved p20 Casp-1 intensity normalized to β-actin. (**G**) Plasma cell-free DNA concentrations in patients with myocardial infarction (MI). (**H**) Representative image of gelatine zymography of plaque lysates from asymptomatic or symptomatic patients (quantification in Suppl. Fig. 11). (**I**) Quantification of F.XIIa in plaque lysates from asymptomatic or symptomatic patients. All statistical tests in this figure: U test.

## Discussion

Early recurrent events after ischemic stroke present a pressing sociomedical problem with an unmet need. Current secondary prevention therapies target mainly blood lipid levels (statins) and platelet aggregation (aspirin), which are effective for long-term prevention of vascular events. However, these therapies have only a negligible effect on the very high recurrence rate after large artery stroke^4^, most likely because this is driven by so far largely unknown immunological mechanisms^11^. Here, we identified inflammasome activation by cell-free DNA as initiator of an inflammatory cascade leading to atherosclerotic plaque degradation and thrombosis. Our results propose the neutralization of cell-free DNA or inhibition of downstream inflammasome activation as highly efficient therapeutic approaches to prevent recurrent vascular events.

The CANTOS trial–testing the use of IL-1β-specific antibody treatment in patients with previous myocardial infarction–clearly highlighted the relevance of residual inflammatory risk and demonstrated the potential of anti-inflammatory therapies to prevent recurrent ischemic events^21^. However, targeting this central cytokine of innate immune defense also significantly increased the risk of fatal infections, which might preclude further clinical development of such unspecific anti-inflammatory strategies^22^. A critical limitation of previous clinical trials targeting inflammatory mechanisms in high-risk atherosclerotic patients was the insufficient mechanistic knowledge on the exact immunological events that resulted in recurrent vascular events. This knowledge gap has so far prevented the development of more specific, and thereby safer, therapeutic strategies targeting molecular mechanisms of stroke recurrence.

Here we identified increased circulating cell-free (cf)DNA concentrations and local activation of the AIM2 inflammasome as the key mechanisms leading to exacerbated plaque inflammasome activation. We demonstrated that the increase in circulating cfDNA was sufficient and causally involved in plaque destabilization using an *in vivo* cfDNA-challenge experiment and validated in human patient samples. Correspondingly, use of recombinant DNase efficiently prevented plaque inflammation, destabilization and recurrent ischemic events. Importantly, no direct immunosuppressive function is known for the experimental or clinical use of DNase in various disease conditions including cystic fibrosis and pleural infections^23–25^. In contrast, we have previously demonstrated that prevention of AIM2 inflammasome activation by DNase treatment might even paradoxically improve immunocompetence during secondary bacterial infections, because reduction of early inflammasome activation in response to tissue injury prevents subacute immunosuppression^18,26^. Of note, lower endogenous DNase activity was recently described as an independent risk factor for stroke-associated infections following severe ischemic stroke^27^. Therefore, the use of DNase is a promising candidate for further clinical development as a therapeutic approach in early secondary prevention that is highly efficient but also safe due to its specific and non-immunosuppressive function.

Not only the immunogenic mediators leading to inflammatory plaque rupture, but also the exact pathways contributing to local plaque destabilization and recurrent atherothrombosis after stroke were so far elusive. Here, we focused on the role of MMP-mediated plaque degradation and subsequent initiation of the contact-dependent coagulation cascade. In correspondence with our observations, increased plaque MMP activity has been previously associated with markers of increased plaque vulnerability^19^. Additionally, increased blood MMP9 activity was previously associated with worse stroke outcome^28^. Interestingly, we observed that inflammasome activation and specifically the release of local IL-1b can dose-dependently increase MMP activity. It is likely that MMP-mediated degradation of the plaque extracellular matrix (ECM) leads to destabilization and exposition of ECM components to the blood circulation^29^. Correspondingly, we found inflammasome-dependent luminal accumulation of F.XIIa at the CCA plaque – a process which we found to be associated with the occurrence of secondary infarctions. F.XII activation at the site of vascular damage has been previously proposed as an initiator mechanism of thrombus formation on ruptured plaques and its pharmacological targeting was recently demonstrated to stabilize atherosclerotic plaques^30^. Therefore, based on our findings, F.XIIa might represent another potential therapeutic target to prevent recurrent ischemic events and warrants further exploration.

In summary, we present in this study a mechanistic explanation for the high rate of early recurrent events after stroke and myocardial infarction in patients with atherosclerosis. Using a novel animal model of post-stroke plaque rupture and secondary infarctions, we identified the immunological mechanisms and validated these in human carotid artery plaque samples. We confirmed the efficient therapeutic targeting of this signaling cascade with DNase administration as a promising therapeutic candidate for further clinical translation.

## Acknowledgement

The authors would like to thank Kerstin Thuß-Silczak for her excellent technical support, Qihui Zhou for support with ultrasound analysis, Ulrike Schillinger and Rong Fang for support in establishing the small-animal MRI analysis and Oliver Söhnlein for his critical input in study design and established the tandem stenosis model. The authors moreover would like to acknowledge Ricarda D. Stauss, Claudia Schrimpf and Mathias Wilhelmi for their support in collecting human carotid plaque samples.

## Author Contributions

JC and SR performed most experiments, analyzed data and contributed to writing the manuscript; GMG provided human carotid endarterectomy samples; MG, AK, TL and SW acquired and analyzed clinical epidemiological data; SZ, CF, XL, OC, JZ, and YA performed experiments and analyzed data; DB, JPG, HS, and MD contributed critical material and techniques for this study; HBS and MD contributed critical input to study design and manuscript writing; AL initiated and coordinated the study, analyzed data and wrote the manuscript.

