## Supplementary figures and methods for "Stroke induces early recurrent vascular events by inflammasome-dependent atherosclerotic plaque rupture"

#### Supplementary material list:

- Supplementary Figures 1-11

Suppl. Fig. 1: Established secondary prevention fails to attenuate early post-stroke vascular inflammation and atheroprogression.
Suppl. Fig. 2: Animal model of rupture-prone CCA plaques
Suppl. Fig. 3: Stroke accelerates plaque destabilization and causes plaque rupture
Suppl. Fig. 4: Myocardial infarction exacerbates plaque vulnerability
Suppl. Fig. 5: Stroke exacerbates cellular plaque inflammation
Suppl. Fig. 6: Stroke induces inflammasome activation in atherosclerotic plaques
Suppl. Fig. 7: Inflammasome inhibition alleviates vascular inflammation and stabilizes atherosclerotic plaques after stroke
Suppl. Fig. 8: Post-stroke plaque inflammasome activation is mediated by cell-free DNA
Suppl. Fig. 9: Stroke increases matrix metalloproteinase activity in atherosclerotic plaques
Suppl. Fig. 10: Stroke initiates the intrinsic coagulation cascade at atherosclerotic plaques
Suppl. Fig. 11: Blood leukocyte counts do not differ between stroke and asymptomatic patients with high-grade atherosclerosis

- Supplementary materials and methods

- Supplementary references

**Supplementary Figure 1**

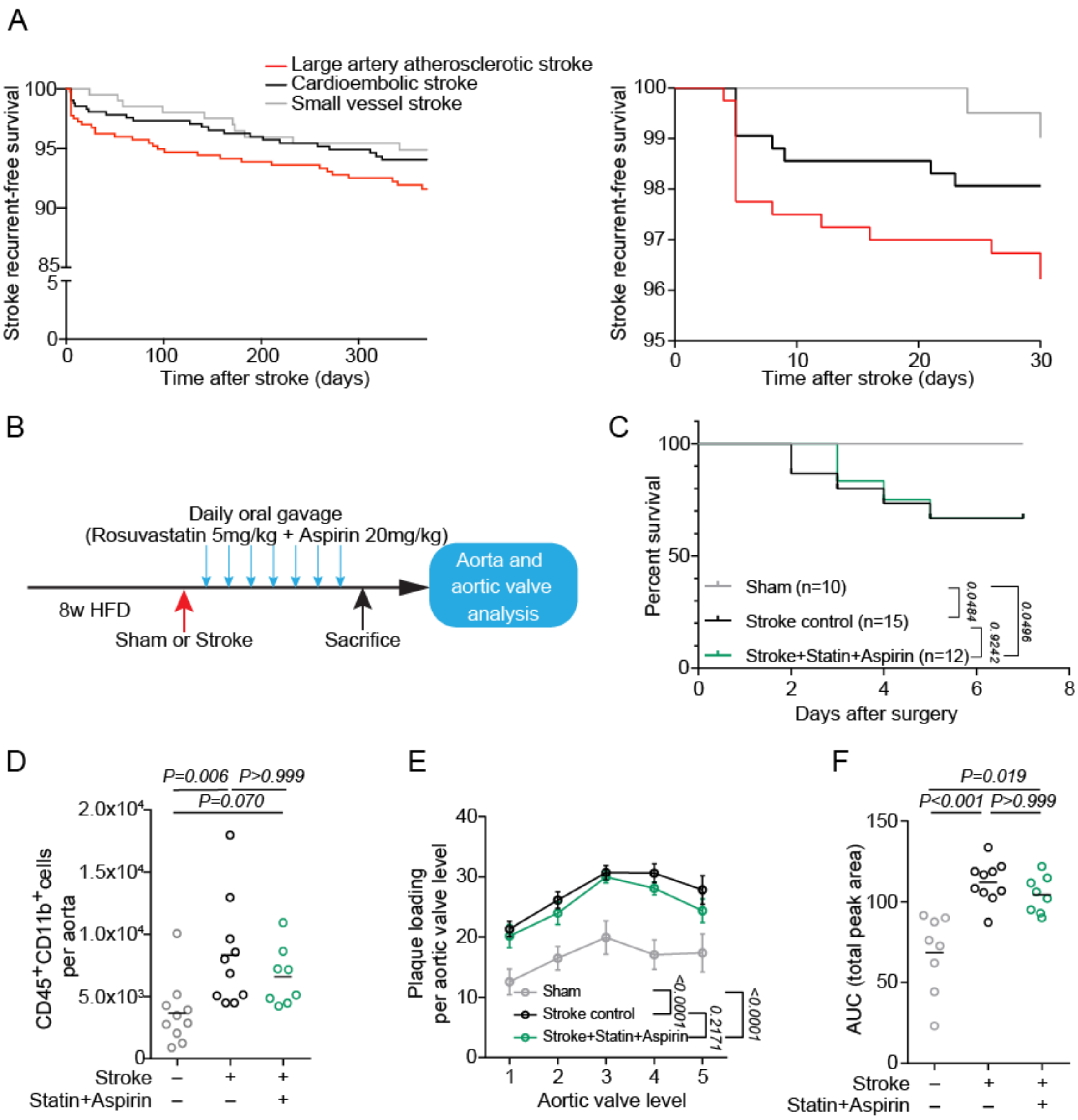


**Supplementary Figure 1. Established secondary prevention fails to attenuate early post-stroke vascular inflammation and atheroprogression.** (**A**) Kaplan-Meier curves for recurrence-free survival of stroke patient from the same cohorts (PROSCIS and DEMDAS/DEDEMAS) as shown in Figure 1A. Recurrence-free survival is illustrated by etiology of the incident stroke for the full time range of 1 year after the incident event (left) and magnified for the first post-stroke month (right). Recurrence risk is highest after large artery atherosclerosis stroke in the early (first week) after the incident event. (**B**) Experimental design: 8 w-old HFD fed *ApoE*$^{-/-}$ mice underwent sham or stroke surgery. Mice were treated orally with either control or a combination of Rosuvastatin (5 mg/kg) and Aspirin (20 mg/kg) for 7 consecutive days after stroke. (**C**) Kaplan-Meier survival curves of stroke control, statin and aspirin treated, or sham operated mice. Mantel-Cox test; n = 10 (sham), n = 15 (control), n = 12 (statin + aspirin treatment). (**D**) Flow cytometry analysis of whole aorta cell suspensions for total monocyte (CD45$^{+}$CD11b$^{+}$) cell counts between control or treated mice after stroke compared to sham-operated mice (ANOVA, n = 8-10 per group). (**E, F**) Quantification of aortic valve plaque load displayed no differences between stroke control and statin + aspirin-treated mice. Data is shown as (D) percentage of plaque area per aortic valve level and (E) area under the curve (ANOVA, n = 8-10 per group).

**Supplementary Figure 2**

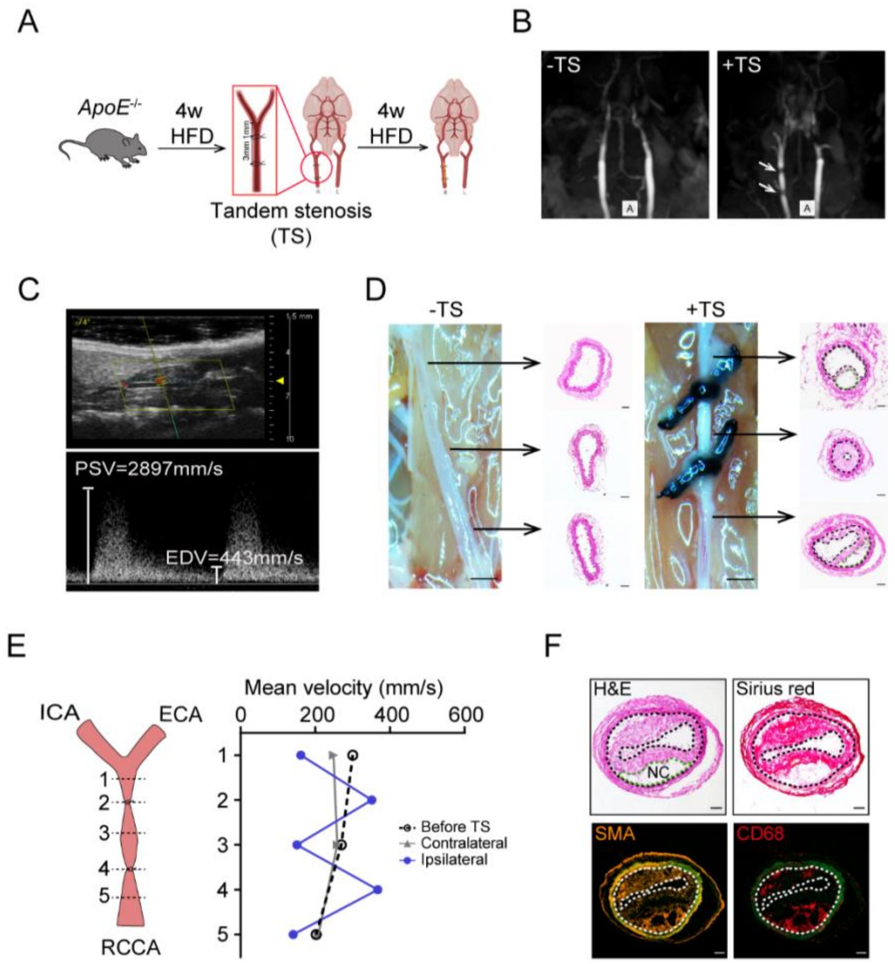


**Supplementary Figure 2. Animal model of rupture-prone CCA plaques** (**A**) Schematic illustration of the tandem stenosis (TS) model for induction of vulnerable atherosclerotic plaques: 8 w-old HFD fed *ApoE*$^{-/-}$ mice received tandem stenosis (TS) surgery on the right common carotid artery (RCCA). Mice were then fed with high fat diet for an additional 4 w. (**B**) Representative images of CCA MRI TOF sequence 4 w after mice received control or TS surgery. White arrows highlight the two ligations on the RCCA. (**C**) Representative pulse-wave mode ultrasound image of the RCCA 4 w after TS, imaged at the location of proximal ligation at 40 MHz (upper panel). Corresponding CCA velocity waveform measured at the location of proximal ligation location 4 w after TS surgery (lower panel). PSV: peak systolic velocity, EDV: end diastolic velocity. (**D**) Representative photograph of the CCA anatomy and corresponding H&E staining for each vessel segment 4 w after TS surgery of both CCAs (-TS represents contralateral (left) CCA without TS ligation; +TS represents ipsilateral (right) CCA with TS ligation, scale bar = 100 µm; H&E staining, scale bar = 50 µm). (**E**) Schematic description of locations for blood flow measurement on the right CCA (ICA: internal carotid artery; ECA: external carotid artery; RCCA: right common carotid artery). Corresponding quantification of mean velocity in the CCA measured at both CCAs before and 4 w after TS surgery (right). (**F**) Representative images of the unstable plaque in the right CCA 4 w after TS surgery (area between two dotted lines indicates intima area, green dotted line indicates necrotic core, scale bar = 50 µm).

**Supplementary Figure 3**

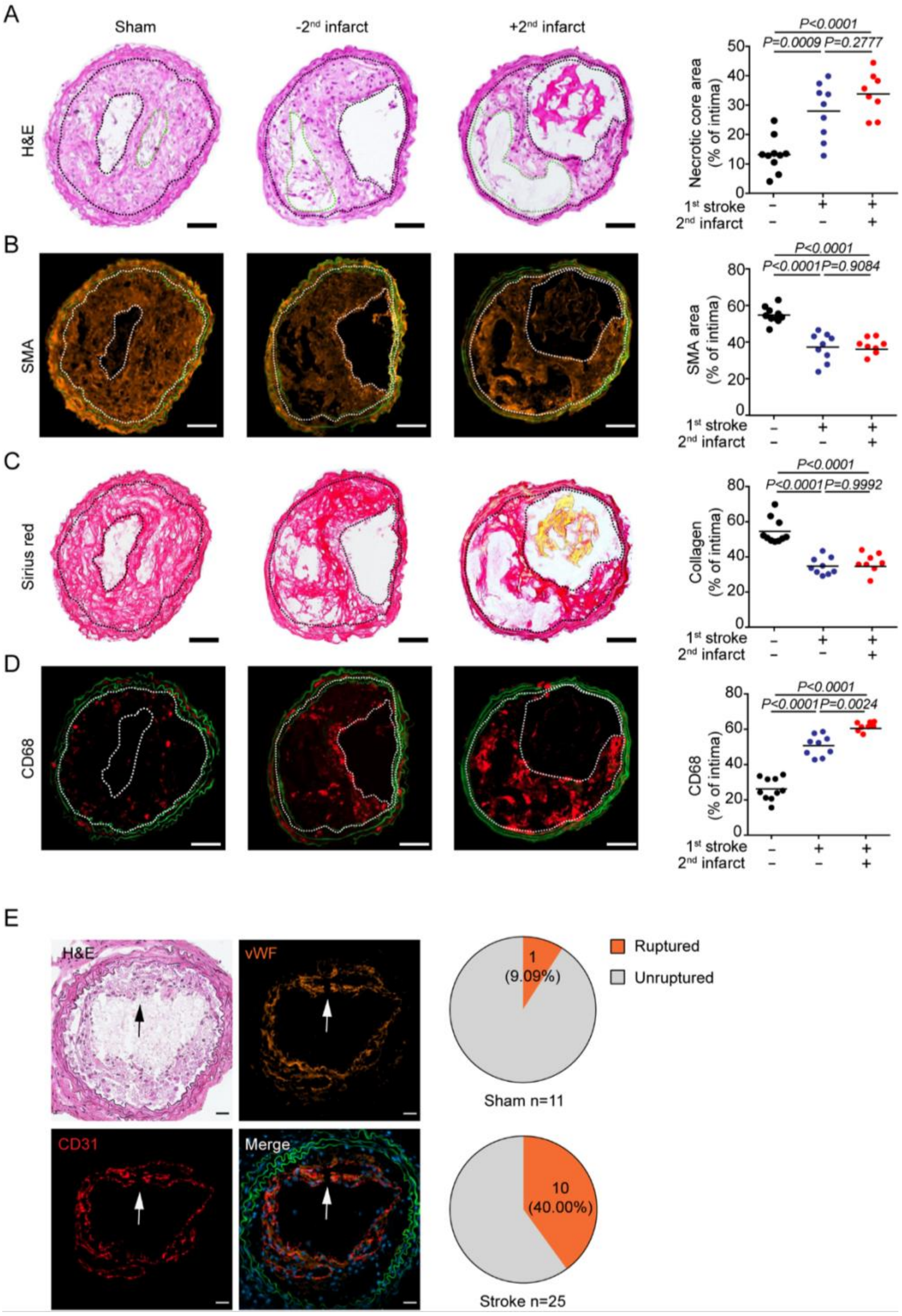


**Supplementary Figure 3. Stroke accelerates plaque destabilization and causes plaque rupture**. (**A**) Representative microphotographs of CCA H&E staining. Area between two black dotted lines represents intima. Green dotted lines represent necrotic cores. Necrotic core area was quantified as the percentage of total intima area (ANOVA, n=8-10 per group). (**B**) Representative images of smooth muscle actin (SMA) immunofluorescence staining. SMA area was quantified as the percentage of total intima area (ANOVA, n=8-10 per group). (**C**) Representative microphotographs of picro sirius red staining. Collagen area was quantified as the percentage of total intima area (ANOVA, n=8-10 per group). (**D**) Representative microphotographs of CD68 immunofluorescence staining . Images were segmented by thresholding to convert fluorescence signal into a binary image. CD68 area was quantified as the percentage of total intima area (ANOVA, n=8-10 per group). (**E**) Arrows indicate a disrupted fibrous cap in lesion. The pie charts illustrates the proportion of mice with ruptured CCA plaques 1 w after sham or stroke surgery (all scale bars = 50 μm).

**Supplementary Figure 4**

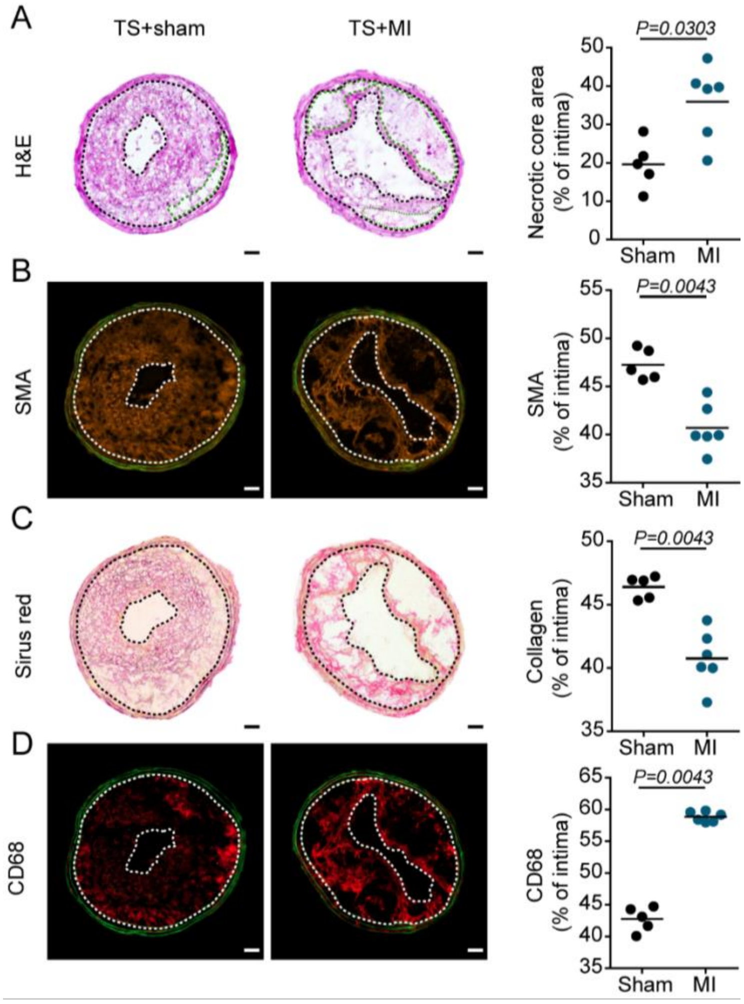


**Supplementary Figure 4. Myocardial infarction exacerbates plaque vulnerability**. Representative microphotographs of H&E (**A**), SMA (**B**), picro sirius red (**C**) and CD68 (**D**) staining in CCA sections 1 w after sham or myocardial infarction (MI) surgery (scale bar = 50 µm). Area between two dotted lines indicate intima area. Green dotted line represents necrotic core area. Corresponding quantification of necrotic core area, SMA, collagen and CD68 area 1 w after sham or MI operated mice (quantification were performed as described in Suppl. Fig. 3, U test, n= 5-6 per group).

**Supplementary Figure 5**

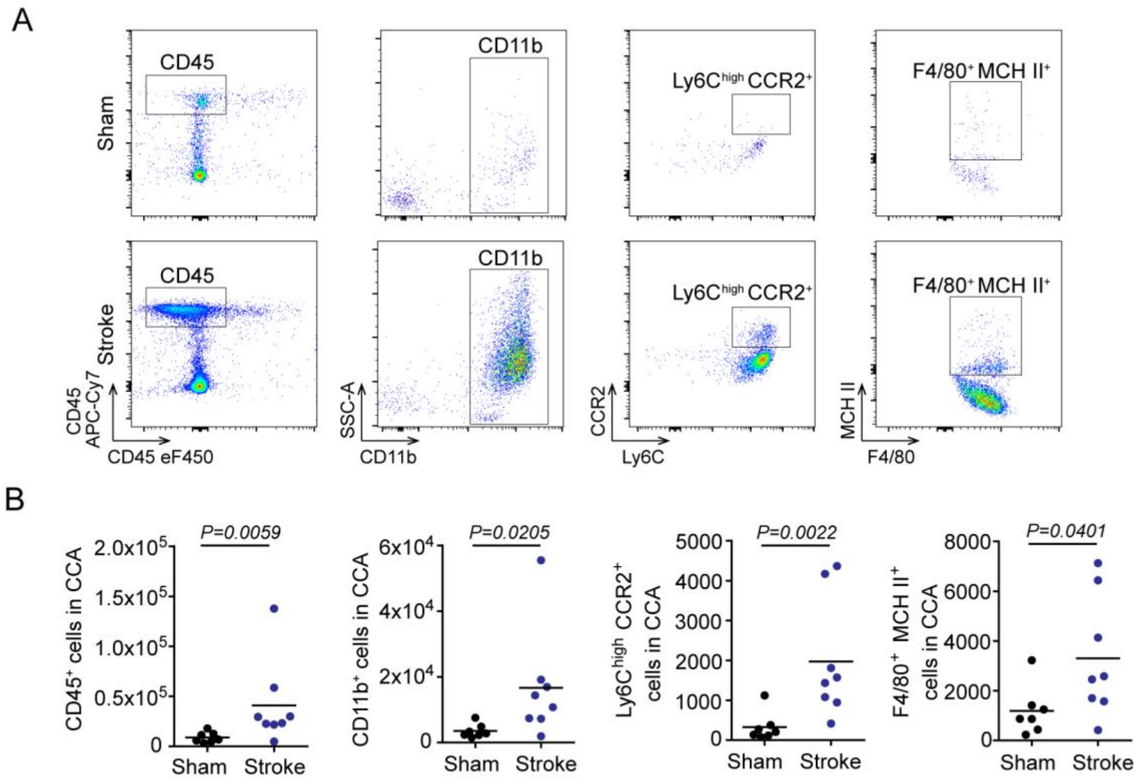


**Supplementary Figure 5. Stroke exacerbates cellular plaque inflammation.** (**A**) Representative gating strategy for flow cytometry analysis of whole CCA cell suspensions 24h after sham or stroke surgery. (**B**) Flow cytometry analysis of CCA cell suspensions showing total leukocytes ($CD45^{+}$), monocytes ($CD11b^{+}$), proinflammatory subset ($Ly6C^{high}$ $CCR2^{+}$) and macrophages ($F4/80^{+}$ $MCHII^{+}$) cell counts after experimental stroke compared to sham (U test, n=7-8 per group).

**Supplementary Figure 6**

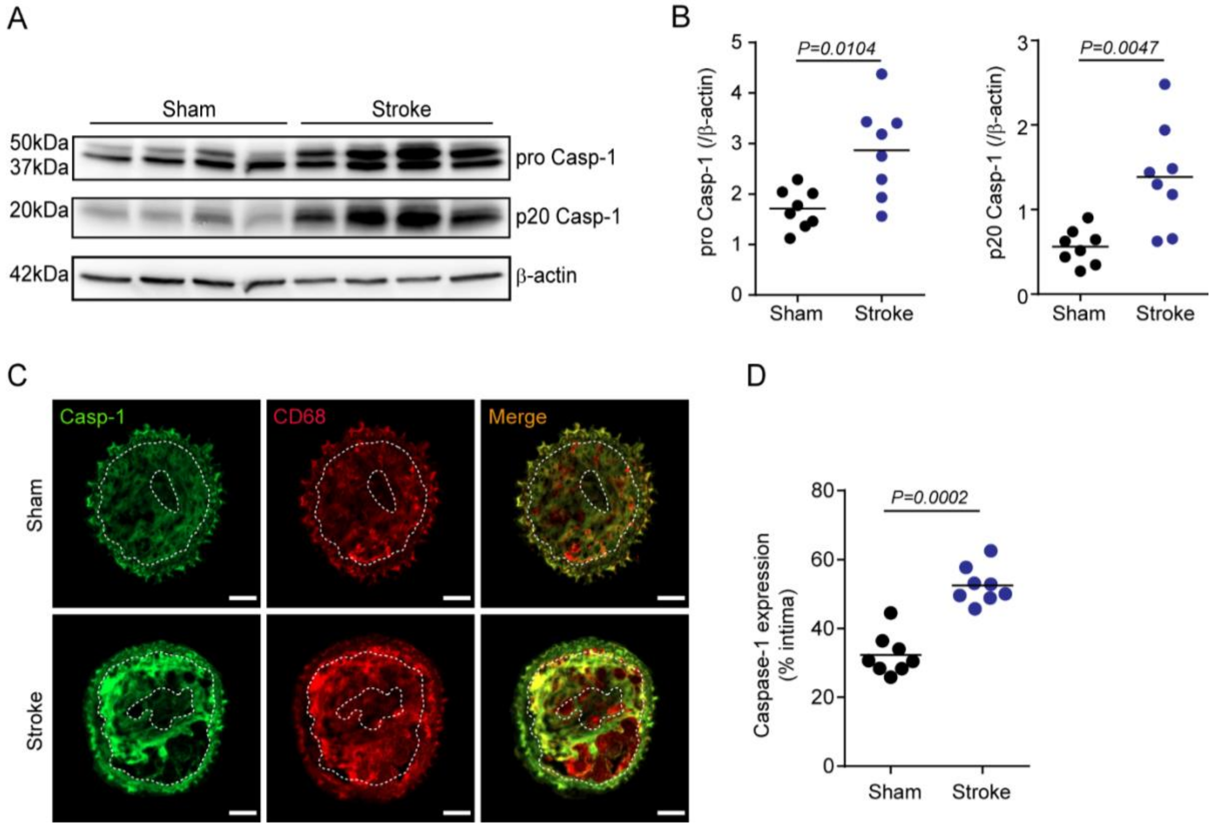


**Supplementary Figure 6. Stroke induces inflammasome activation in atherosclerotic plaques**. (**A**) Representative immunoblot of different cleavage forms of caspase-1 in CCA lysates with TS 1 w after sham or stroke surgery. (**B**) Quantification of pro Caspase-1 (ProCasp-1) and cleaved caspase-1 (p20 Casp-1) intensity normalized to β-actin (U test, n=8 per group). (**C**) Representative immunofluorescence staining of caspase-1 (Casp-1) in CCA sections 1 w after sham or stroke surgery (scale bar = 50 µm). Images were segmented by thresholding to convert fluorescence signal into a binary image. Area between two white dotted lines represent intima. (**D**) Caspase-1 expression was quantified as the percentage of total intima area (U test, n=8 per group).

**Supplementary Figure 7**

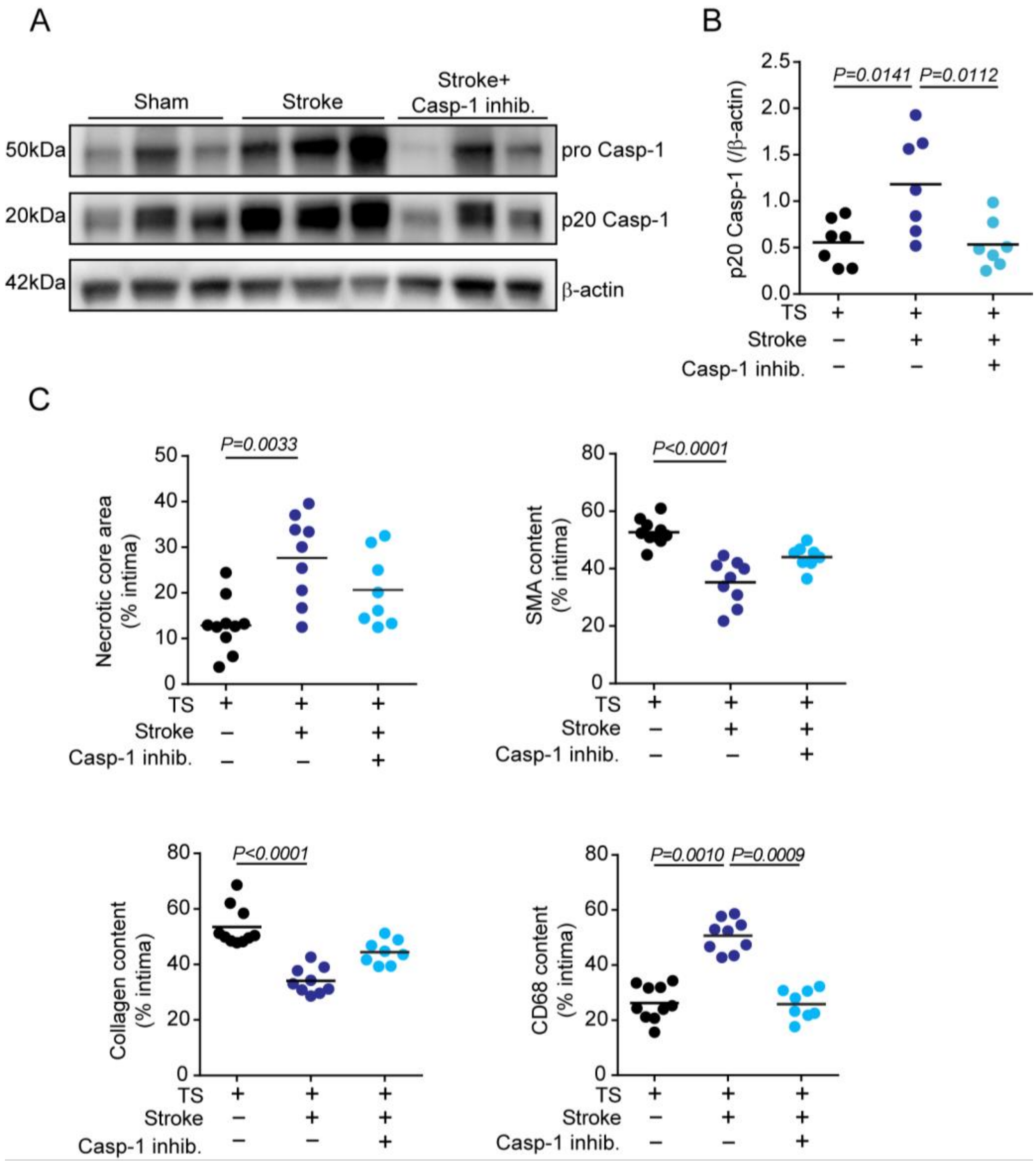


**Supplementary Figure 7. Inflammasome inhibition alleviates vascular inflammation and stabilizes atherosclerotic plaques after stroke.** (**A**) Representative immunoblot image of the different cleavage forms of caspase-1 (Casp-1) in CCA lysates 1 w after sham, stroke control and stroke + caspase-1 inhibitor (VX 765) administration. (**B**) Quantification of cleaved p20 Casp-1 intensity normalized to β-actin in CCA lysates (+TS) 1 w in the three treatment groups (black: sham; blue: stroke; light blue: stroke + VX765, ANOVA, n= 7 per group). (**C**) Quantification of necrotic core area, SMA, collagen and CD68 area 1 w after sham or stroke in the respective treatment groups (performed as shown in Suppl. Fig. 3, ANOVA, n=8-10 per group).

### Supplementary Figure 8

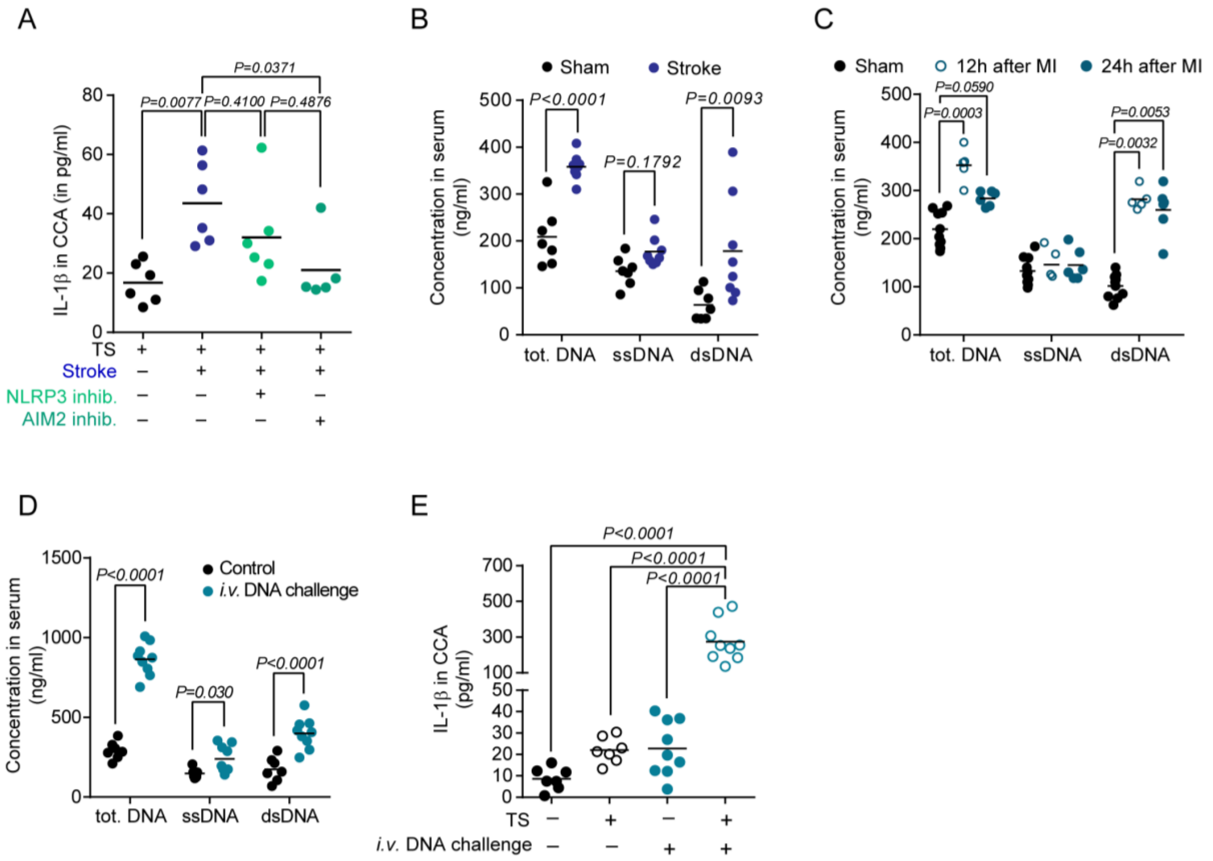


**Supplementary Figure 8. Post-stroke plaque inflammasome activation is mediated by cell-free DNA**. (**A**) ELISA analysis of IL-1β in CCA lysates from mice with tandem stenosis (TS), 24 hours after stroke in control-, NLRP3 inhibitor- (MCC950) or AIM2 inhibitor- (4-sulfonic calix[6]arene) treated mice, and in sham operated mice (ANOVA, = 5-6 per group). (**B**) Total cell-free DNA (cfDNA), single-stranded DNA (ssDNA) and double-stranded DNA (dsDNA) in mouse serum 24h after sham or stroke surgery (U test, n=7-8 per group). (**C**) Total cell-free DNA (tot.DNA), single-stranded DNA (ssDNA) and double-stranded DNA (dsDNA) in mice serum after sham or 12 h, 24 h after myocardial infarction (MI) surgery. (**D**) tot.DNA, ssDNA and dsDNA in mice serum were measured 24 h after control or *i.v.* DNA challenge (multiple t test, control, n = 7; DNA challenge, n = 9). (**E**) ELISA analysis for IL-1β in CCA lysates 24 hours after *i.v.* DNA challenge (ANOVA, n =7-9 per group).

**Supplementary Figure 9**

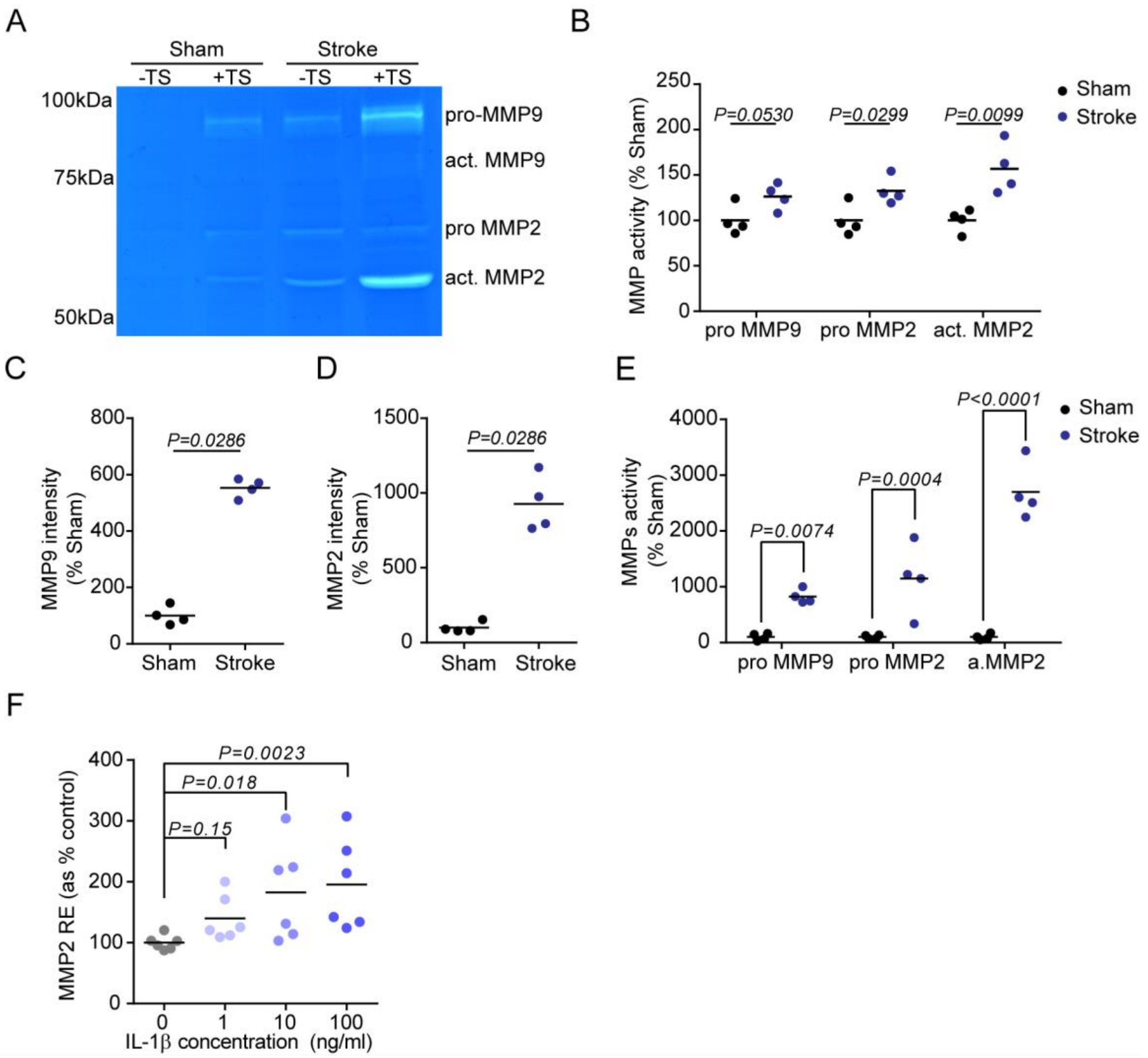


**Supplementary Figure 9. Stroke increases matrix metalloproteinase activity in atherosclerotic plaques.** (**A**) Representative images of gelatin zymography of CCA lysates for MMP activity in mice 1 week after sham or stroke surgery. The region of MMP activity appears as a clear band against dark blue background where the MMP has digested the gelatin substrate on the zymogram gel. (**B**) MMP activity shown in (A) was quantified as the gelatin digestion area 1 w after stroke surgery normalized to sham operated mice (multiple t test, n=4 per group). (**C, D**) Quantification of MMP9 and MMP2 intensity from immunoblot micrograph shown in Fig. 3B (normalized to sham stimulated, U test, n= 4 per group). (**E**) MMP activity shown in Fig. 3B was quantified as the gelatin digestion area in the stroke serum-stimulated medium normalized to sham serum-stimulated group (t test, n= 4 per group). (**F**) Relative expression (RE) of MMP2 expression in WT BMDMs after IL-1β stimulation was quantified as the percentage of the control group (H test, n= 6 per group).

**Supplementary Figure 10**

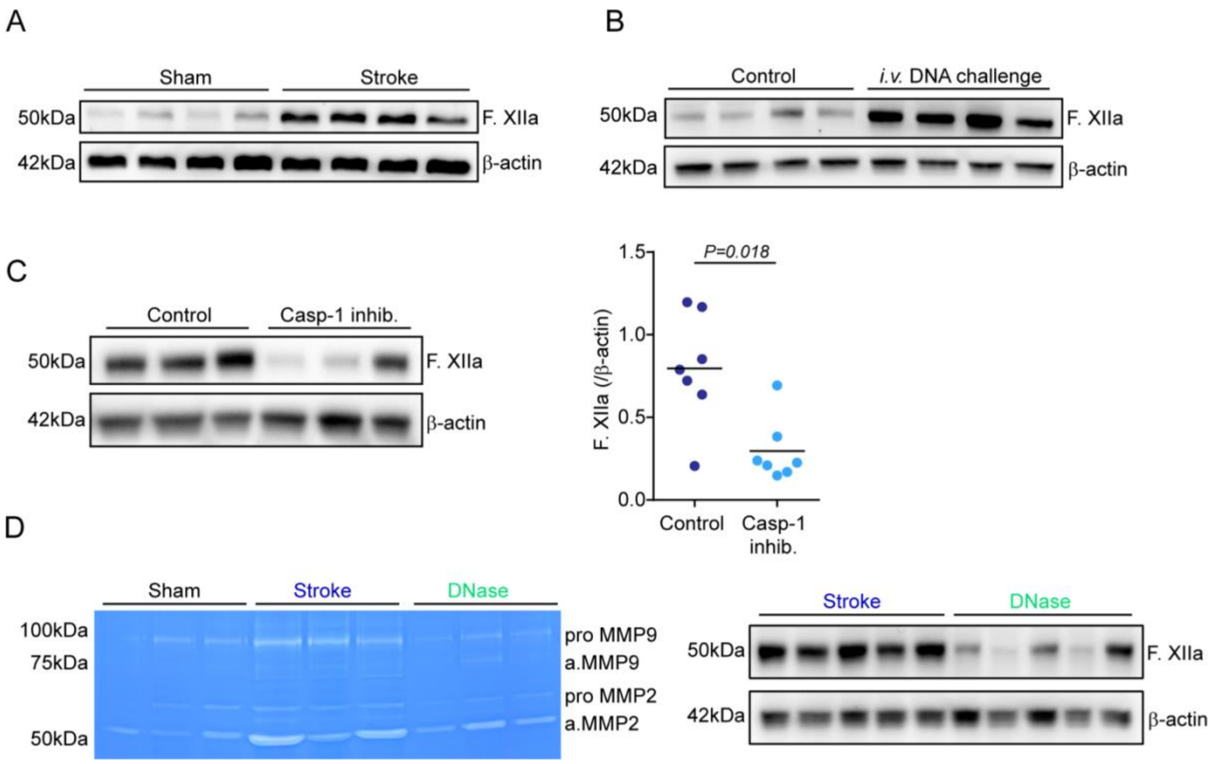


**Supplementary Figure 10. Stroke initiates the intrinsic coagulation cascade at atherosclerotic plaques**. All analyses were performed on CCA lysates in mice with stenotic CCA plaques after TS surgery in HFD fed *ApoE*$^{-/-}$. (**A**) Representative immunoblot of activated Factor XII (F.XIIa) 1 w after stroke or sham surgery. (**B**) Representative immunoblot micrograph of FXIIa in CCA lysates 24 hours after *i.v.* DNA challenge. (**C**) Representative immunoblot of the F.XIIa in CCA lysates 24 h after stroke in mice treated with control treatment or caspase-1 inhibition (VX765). Corresponding quantification of F.XIIa intensity normalized to β-actin in CCA lysates 1 w after stroke in control- or caspase-1 inhibitor- treated mice (U test, n = 7 per group). (**D**) Representative gelatin zymography of CCA lysates for MMP activity in mice 24 h after sham, stroke or stroke + DNase treatment (left). Representative immunoblot of F.XIIa in +TS CCA lysates 24h after surgery (right).

**Supplementary Figure 11**

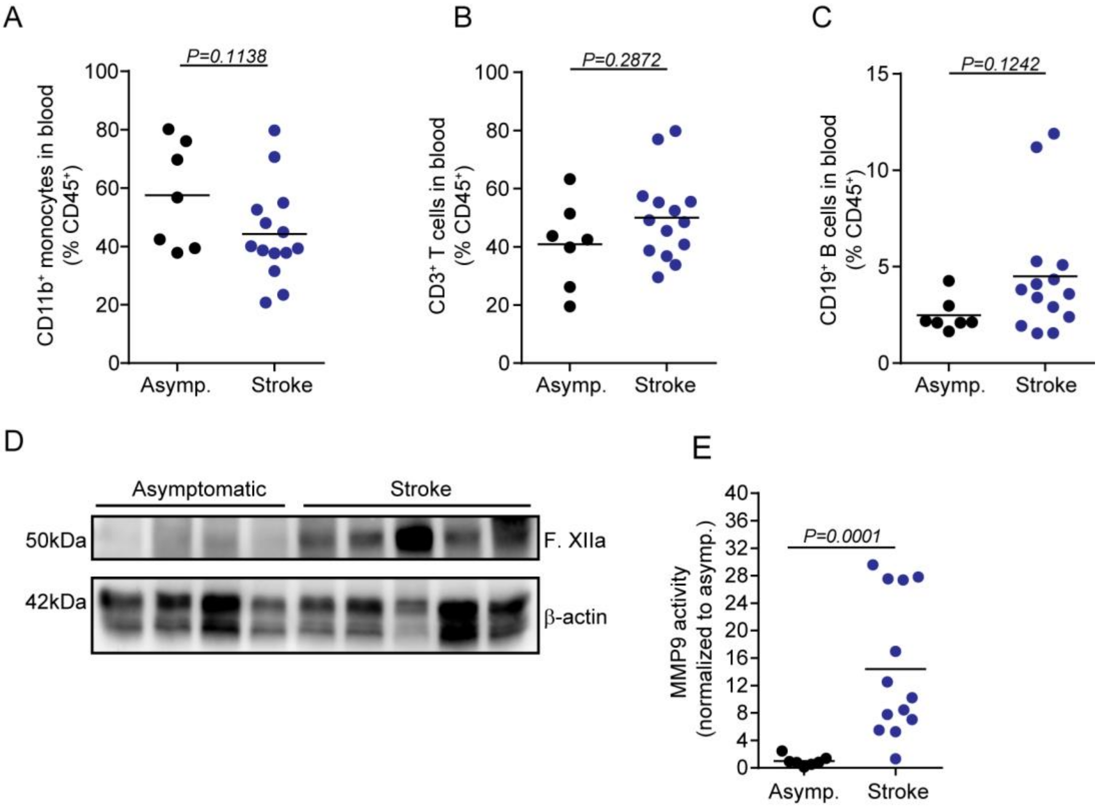


**Supplementary Figure 11. Blood leukocyte counts do not differ between stroke and asymptomatic patients with high-grade atherosclerosis.** (**A-C**) Flow cytometry analysis of blood from asymptomatic patients or stroke patients showing the percentage of monocytes ($CD11b^+$), T cells ($CD3^+$) and B cells ($CD19^+$) out of total leukocytes ($CD45^+$) (U test, asymptomatic patients, n = 7; symptomatic patients, n = 13). (**D**) Representative immunoblot from asymptomatic and stroke patients for F.XIIa and β-actin (Quantification can be found in Figure 4I). (**E**) Quantification of MMP9 activity normalized to the activity in asymptomatic patients (Representative image shown in Figure 4H).

### Supplementary Materials and Methods

#### Animal experiments

All animal experiments were performed in accordance with the guidelines for the use of experimental animals and were approved by the government committee of Upper Bavaria (Regierungspraesidium Oberbayern; #02-2018-12). *ApoE*$^{-/-}$ (B6.129P2-Apoetm1Unc/J; JAX strain: 002052) were bred and housed at the animal core facility of the Centre for Stroke and Dementia Research (Munich, Germany). *ApoE*$^{-/-}$ mice were fed an HFD (#88137, ssniff) from 8 weeks on.

For this exploratory study, animal numbers were estimated based on previous results from the transient ischemia-reperfusion stroke model on extent and variability of atheroprogression after stroke. Data were excluded from all mice that died during surgery. Detailed exclusion criteria are described below. Animals were randomly assigned to treatment groups and all analyses were performed by investigators blinded to group allocation. All animal experiments were performed and reported in accordance with the Animal Research: Reporting of In Vivo Experiments (ARRIVE) guidelines[1].

#### Drug Administrations

Oral gavage with Aspirin and Rosuvasatin

Mice received a daily bolus of Aspirin (20mg kg$^{-1}$, Sigma Aldrich, Germany) and Rosuvastatin (5mg kg$^{-1}$, Sigma Aldrich, Germany) via oral gavage. Aspirin and Rosuvastatin were solved in water (sterile dd$H_2O$) and mixed with powdered chow diet (ssniff). A single daily bolus was 500 µl.

Recombinant DNase 1 (DNase 1)

*ApoE*$^{-/-}$ mice received DNase injections as we previously described[2]. Briefly, 1000 U recombinant DNase 1 (Roche, Switzerland) dissolved in incubation 1x buffer (40 mM Tris-HCl, 10 mM NaCl, 6 mM MgCl2, 1 mM CaCl2, pH 7.9, diluted in PBS, Roche) was injected *i.v.* via tail vein right before surgery in a final volume of 100 µl. The control group was administered saline injections at the same volume, routine, and timing as experimental group.

Caspase-1 inhibitor (VX-765)

The caspase-1 inhibitor VX-765 (stock in DMSO) was dissolved in PBS (Belnacasan, Invivogen, US) and injected *i.p.* 1 h prior to surgery at a dose of 100 mg kg$^{-1}$ body weight at a final volume of 300 µl[2]. The control group was administrated saline injections at the same volume, routine, and timing as experimental group.

NLRP3 inflammasome inhibitor (MCC-950)

Mice received two injections (1 h prior to and 1 h after surgery) of the NLRP3 inflammasome inhibitor MCC-950 dissolved in sterile saline at a dose of 50mg kg$^{-1}$ body weight (Invivogen, US). MCC-950 or the control (sterile saline) was injected *i.p.* in a final volume of 200 µl.

AIM2 inhibitor (4-sulfonic calix[6]arene)

The AIM2 inhibitor 4-sulfonic calix[6]arene was recently characterized by Green et al. [3]. The stock solutions (in DMSO) was dissolved in PBS and injected *i.p.* 1 h prior to surgery at a dose of 5 mg kg$^{-1}$ body weight at a final volume of 200 µl. The control group was administered control injections (sterile saline) at the same volume, routine, and timing.

DNA challenge

DNA derived from stimulated neutrophils (see below for stimulation and isolation of NET DNA) was injected *i.v.* at a dose of 5 µg at a final volume of 200 µl (sterile saline). The control group was administrated control injections (sterile saline) at the same volume, routine, and timing.

**Patient cohorts for epidemiological analysis**

The analysis presented in Figure 1A was performed using patient data from two multi-center prospective observational hospital-based cohort studies of patients with acute ischemic stroke in Germany (PROSCIS and DEMDAS/DEDEMAS). The study protocols and the detailed baseline patient characteristics have been described before[4,5]. Basic demographic and stroke-related characteristics are summarized below. Briefly, for both the PROSCIS and the DEMDAS/DEDEMAS cohorts, patients ≥18 years of age with an acute ischemic stroke confirmed with neuroimaging and with symptom onset in the last 7 and 5 days, respectively, were recruited through the local stroke units of seven tertiary stroke centers in Germany. Patients in PROSCIS underwent follow-up telephone interviews at 3 and 12 months after stroke, whereas patients in DEMDAS/DEDEMAS underwent a telephone interview at 3 months after stroke as well as face-to-face interviews and inspection of their medical records by a physician at 6 and 12 months after stroke. The outcome of interest for the current analysis included the occurrence of a recurrent ischemic stroke or transient ischemic attack within the first 12 months after stroke, as self-reported by the patient and documented in their medical records.

| | PROSCIS | DEMDAS/DEDEMAS |
|---|---|---|
| Sample size (n) | 1083 | 715* |
| Age (y), mean (SD) | 67.6 (13.4) | 68.0 (11.2) |
| Sex (% males) | 59.7 | 66.7 |
| TOAST subtype (%) | | |
| Large artery atherosclerosis | 20.9 | 26 |
| Cardioembolism | 24.1 | 23.4 |
| Small artery occlusion | 11.9 | 11.7 |
| Other etiology | 3.5 | 4.2 |
| Unknown etiology | 39.6 | 34.7 |

* 21 patients with hemorrhagic stroke originally included in the DEMDAS/DEDEMAS study have been excluded from the current analysis.

**Patient cohort for carotid endarterectomy sample analysis**

Patients scheduled for carotid endarterectomy due to symptomatic or asymptomatic carotid stenosis were prospectively recruited at the Department of Neurology and Cardiothoracic-Transplantation- and Vascular Surgery at Hannover Medical School between June 2018 and December 2020. Carotid stenosis was defined as symptomatic if cerebral ischemia occurred in the territory of the affected artery and concurrent stroke etiologies were excluded following standardized stroke diagnostics including cranial computed tomography (CT) and/or magnetic resonance (MR) imaging. CT or MR-angiography, transthoracic or transesophageal echocardiography, cardiac rhythm monitoring and Doppler/duplex ultrasound. Peripheral venous blood was drawn immediately prior to surgery and EDTA whole blood samples were used for flow cytometry analysis. Carotid plaque samples were obtained during carotid endarterectomy and immediately preserved in phosphate-buffered saline. Both blood and tissue samples were sent for further analysis on the same day of collection. All patients providde written informed consent and the ethics committee at Hannover Medical School approved the study.

Thirteen patients with symptomatic and seven patients with asymptomatic, high-grade carotid stenosis were recruited. Median age was 73 years ($25^{th}$-$75^{th}$ percentile: 62-80 years). See supplemental table for clinical and demographic patient details.

| | Symptomatic carotid stenosis (n=13) | Asymptomatic carotid stenosis (n=7) |
|---|---|---|
| Age (median (a); $25^{th}$-$75^{th}$ percentile) | 64 (61-75) | 78 (76-82) |
| Sex (male, n) | 12 (92%) | 6 (86%) |
| Arterial hypertension (n) | 11 (84%) | 7 (100%) |
| Diabetes mellitus (n) | 4 (31%) | 3 (43%) |
| BMI (median (kg/m²); $25^{th}$-$75^{th}$ percentile) | 25.5 (23.2-29.3) | 26.8 (24.5-31.6) |
| Current nicotine consumption (n) | 5 (38%) | 0 (0%) |
| Dyslipidemia (n) | 9 (69%) | 6 (85%) |
| Coronary artery disease | 3 (23%) | 3 (43%) |
| Peripheral artery disease | 1 (8%) | 2 (29%) |
| History of myocardial infarction | 1 (8%) | 1 (14%) |
| History of stroke | 2 (15%) | 0 (0%) |

**Patient cohort for myocardial infarction (MI) sample analysis**

STEMI patients were prospectively recruited between September 2016 and February 2018 at the German Heart Centre Munich and the Klinikum rechts der Isar (both at the Technical University of Munich).

The diagnosis of STEMI was based on chest pain within the last 12 hours, persistent ST-segment elevation ≥ 1 mm in at least 2 extremities or ≥ 2 mm in at least 2 chest leads and diagnosis of type 1 myocardial infarction according to cardiac catheterization. Exclusion criteria were: cardiogenic shock, LV-EF ≤ 35, co-existing chronic or inflammatory diseases, anti-inflammatory drug therapy (e.g. cortisol), myocardial infarction type 2 – 5. Blood samples for plasma analysis were collected in EDTA tubes immediately after admission to the hospital or latest 6 hours after coronary intervention.

Age- and sex-matched patients with known stable coronary artery disease served as controls. They were prospectively recruited between February 2017 and February 2018 during consultation in the outpatient department of the German Heart Centre Munich for routine examination. Exclusion criteria were: history of myocardial infarction, reduced LV-EF, chronic or inflammatory diseases, anti-inflammatory drug therapy. Blood samples for plasma analysis were collected in EDTA tubes on the day of consultation in the outpatient department. All patients provided written informed consent and the institutional ethics committee at Technical University Munich approved the study (235/16 S). EDTA tubes of both STEMI and control patients were centrifuged at 4°C and 2000xg for 15 minutes directly after collection. Plasma aliquots were stored at -80°C for further analysis.

| | Control (n=10) | STEMI (n=10) |
|---|---|---|
| Age (median (a); $25^{th}$-$75^{th}$ percentile) | 65 (52-75) | 52 (48-60) |

| | | |
|---|---|---|
| Sex (male, n) | 5 (50%) | 7 (70%) |
| Arterial hypertension (n) | 9 (90%) | 7 (70%) |
| Diabetes mellitus (n) | 4 (40%) | 0 (0%) |
| BMI (median (kg/m²); 25th-75th percentile) | 24.7 (23.5-28.7) | 26.75 (23.2-28.1) |
| Current nicotine consumption (n) | 2 (20%) | 6 (60%) |
| Dyslipidemia (n) | 9 (90%) | 6 (60%) |
| Coronary artery disease | 10 (100%) | 10 (100%) |
| Peripheral artery disease | 2 (20%) | 4 (40%) |
| History of myocardial infarction | 0 (0%) | 1 (10%) |
| History of stroke | 1 (10%) | 0 (10%) |

**Carotid tandem stenosis model**

Tandem stenosis (TS) surgery was performed as previously described by Chen et al.[6]. Briefly, at 12 weeks of age, 4 weeks after commencement of HFD, *ApoE*$^{-/-}$ mice (C57BL/6J background) were anesthetized with 2 % isoflurane delivered in a mixture of 30 % and 70 % $N_2O$. An incision was made in the neck and the right common carotid artery was dissected from circumferential connective tissues. To control the stenosis diameter, a 150 µm or 450 µm pin was placed on top of the exposed right common carotid artery, with the distal point 1 mm away from the bifurcation and proximal point 3 mm from the distal stenosis, subsequently, a 6-0 braided polyester fibre suture was tied around both the artery and needle, and then the pin was carefully removed. Animals were fed with HFD for additional 4 weeks after TS surgery.

**Ischemia-Reperfusion Stroke Model**

4 weeks after TS surgery, *ApoE*$^{-/-}$ mice were anaesthetized with 2 % isoflurane delivered in a mixture of 30 % $O_2$ and 70 % $N_2O$. An incision was made between the ear and the eye in order to expose the temporal bone. Mice were placed in supine position, and a laser Doppler probe was attached to the skull above the middle cerebral artery (MCA) territory. The common carotid artery and left external carotid artery were exposed via middle incision and further isolated and ligated. A 2-mm silicon-coated filament (Doccol) was inserted into the internal carotid artery, advanced gently to the MCA until resistance was felt, and occlusion was confirmed by a corresponding decrease in blood flow (i.e., a decrease in the laser Doppler flow signal by ≥ 80 %). After 60 minutes of occlusion, the animals were re-anesthetized, and the filament was removed. After recovery, the mice were kept in their home cage with ad libitum access to water and food. Sham-operated mice received the same surgical procedure, but the filament was removed in lieu of being advanced to the MCA. Body temperature was maintained at 37 °C throughout surgery in all mice via feedback-controlled heating pad. Exclusion criteria: 1. Insufficient MCA occlusion (a reduction in blood flow to > 20 % of the baseline value). 2. Death during the surgery. 3. Lack of brain ischemia as quantified post-mortem by histological analysis.

**Myocardial Infarction (MI)**

MI surgery was performed as previously described by Hettwer et al.[7]. Briefly, mice were intubated under MMF anaesthesia (midazolam 5.0 mg/kg BW; medetomidine hydrochloride 1.0 mg/kg BW; fentanyl citrate 0.05 mg/kg

BW; intraperitoneally) and thoracotomy was performed in the left intercostal space. The left anterior descending coronary artery was identified and MI was induced by permanent ligation with an 8-0 prolene suture. Atipamezole hydrochloride (5 mg/kg BW) and flumazenil (0.1 mg/kg BW) (AF) injected subcutaneously were used to antagonize MMF anaesthesia. Mice received subcutaneous buprenorphine (0.3 mg/kg BW) as an analgesic every 8 h for 3 days starting at the end of the surgical procedure.

**Ultrasound imaging (mouse carotid artery doppler analysis)**

Carotid artery blood flow in *ApoE*$^{-/-}$ mice was measured with a high-frequency ultrasound imaging system (Vevo 3100LT, VisualSonics, Toronto, ON, Canada) with a 40 MHz linear array transducer (MX550D, VisualSonics, Canada) before and right after TS surgery, and weekly for 4 weeks afterwards. Mice were anesthetized with isoflurane delivered in a mixture of 30 % $O_2$ and 70 % $N_2O$. B-mode, Color-Doppler mode and pulsed Doppler velocity spectrum were recorded from both sides of CCA. For the right common carotid artery (RCCA), five locations were examined: before proximal ligation, near proximal ligation, between two ligations, near distal ligation and above the distal ligations. For the left common carotid artery (LCCA), since it was not ligated, only three locations were measured: proximal, middle and distal part of LCCA. Pulsed Doppler velocity was determined with the sample volume calibrated to cover the entire vascular lumen and the smallest possible angle of interception (< 60°) between the flow direction and the ultrasound beam. The peak systolic velocity (PSV), end diastolic velocity (EDV) were recorded from CCAs of both sides. VevoLab v3.2.0 software was used for ultrasound imaging analysis. The mean velocity (MV) was calculated as: MV=(PSV+2*EDV)/3.

**Magnetic resonance imaging (MRI)**

MRI was performed in a small animal scanner (3T nanoScan® PET/MR, Mediso, with 25 mm internal diameter quadrature mouse head coil) at 2 and 7 days after sham or stroke surgery. For scanning, mice were anesthetized with 1.2% isoflurane in 30 % oxygen/70 % air applied via face mask. Respiratory rate and body temperature (37 +/- 0,5 °C) were continuously monitored via an abdominal pressure sensitive pad and rectal probe and anaesthesia adjusted to keep them in a physiological range. The following sequences were obtained: coronal T2-weighted imaging (2D fast-spin echo (FSE), TR/TE = 3000/57.1 ms, averages 14, resolution 167 x 100 x 500 $\mu m^3$), coronal T1-weighted imaging (2D fast-spin echo (FSE), TR/TE = 610/28,6 ms, averages 14, resolution 167 x 100 x 500 $\mu m^3$), and DWI (2D spin echo (SE), TR/TE = 1439/50 ms , averages 4, resolution 167 x 100 x 700 $\mu m^3$). MEI images then post-processed using Image J.

**Organ and tissue processing**

Mice were deeply anaesthetized with ketamine (120 mg $kg^{-1}$) and xylazine (16 mg $kg^{-1}$) and venous blood was drawn via cardiac puncture of the right ventricle in 50 mM EDTA (Sigma-Aldrich, Germany); the plasma was isolated by centrifugation at 3,000 g for 15 min and stored at -80 °C until further use. Immediately following cardiac puncture, mice were transcardially perfused with ice cold saline. Subsequently, the common carotid arteries (CCA) from both sides as well as the aortic arches and hearts were carefully isolated and embedded in compound (O.C.T., Tissue-tek, Japan), frozen over dry ice and stored at -80 °C until sectioning.

Heads were cut just above the shoulders. Skin was removed from the head and the muscle was stripped from the bone. After removal of the mandibles and the skull rostral to maxillae, whole brain with skull was post-fixed by 4 % paraformaldehyde (PFA) overnight at 4 °C. Subsequently, samples were transferred to a decalcification solution of 0.3 M EDTA (C. Roth, 292.94 g/mol) at pH 7.4 and stored at 4 °C. EDTA solution was changed after 3 days. Samples were immersed with 30 % sucrose in PBS and then frozen down in -20 °C isopentane (Sigma Aldrich, Germany). 15 µm thick coronal sections were obtained at the level of anterior commissure for immunohistochemical analysis. Sections were mounted on SuperfrostPlus Slides (Thermo Scientific, US) and stored in -80 °C.

**Histology and immunofluorescence**

Carotid (5 µm) cryosections were histologically stained with haematoxylin and eosin (H&E) in 100 µm intervals. Total collagen content was assessed by Picrosirius Red staining (Abcam, US) in consecutive sections. For immunofluorescence staining, cryosections were fixed with 4 % PFA followed by antigen blockade using 2 % goat serum blocking buffer containing 1 % Bovine Serum Album (BSA, Sigma), 0.1 % cold fish skin gelatin (Sigma Aldrich, Germany), 0.1 % Triton X-100 (Sigma) and 0.05 % Tween 20 (Sigma). Next, sections were incubated overnight at 4 °C with the following primary antibodies: Rat anti-mouse CD68 (1:200, ab53444, Abcam), mouse anti-mouse smooth muscle actin (1:200, ab7817, Abcam, US), rabbit anti-mouse Ki67 (1:200, 9129S, Cell Signaling, US), mouse anti-mouse caspase-1 (1:200, AG-20B-0042-C100, Adipogen), sheep anti- Von Willebrand Factor (1:100, ab11713, Abcam, US), rat anti-CD31 (1:200, BM4086, OriGene, US). After washing, sections were incubated with secondary antibodies as following: AF647 goat anti-rat (1:200, Invitrogen, US), Cy3 goat anti-mouse IgG H&L (1:200, Abcam, US), AF594 goat anti-rabbit (1:200, Invitrogen, US), AF488 goat anti-mouse (1:200, invitrogen, US), AF594 donkey anti-sheep (1:200, invitrogen, US). Counterstain to visualize nuclei was performed by incubating with DAPI (1:5000, invitrogen, US). Finally, sections were mounted with fluoromount medium (Sigma Aldrich, Germany). Microphotographs of Immunofluorescent samples were taken with confocal microscope (LSM 880, LSM 980; Carl Zeiss, Germany). Histological sections were imaged with an epifluorescent microscope (Axio Imager M2; Carl Zeiss, Germany) and quantified by using ImageJ software (National Institutes of Health, US).

For the visualization of suspected secondary infarct lesions in the contralateral hemisphere, brain sections (15 µm) were first stained for Fluoro Jade C (FJC) to identify degenerating neurons. FJC staining was performed using the "Fluoro-Jade C Ready-to-Dilute" Staining Kit (Biosensis, TR-100-FJ, US) according to the manufacturer's instructions. To confirm the secondary lesion, double staining of microglia marker Iba-1 (1:200, FUJIFILM Wako Pure Chemical Corporation, US) and terminal deoxynuceotidyl transferase dUTP nick end labeling (TUNEL) staining was performed using Click-iT™ Plus TUNEL Assay for In Situ Apoptosis Detection (Alexa Fluor™ 647 dye, Thermo Scientific, C10619) according to the manufacturer's instructions. Brain samples were photographed on an epifluorescence microscope (Zeiss Axiovert 200M, C. Zeiss, Germany) and a confocal microscope (LSM880, C. Zeiss, Germany).

**Mouse CCA plaque analysis**

Plaque vulnerability was assessed as previously described by Silvestre-Roig et al.[8]. In brief, intima, media and necrotic core area was analyzed in H&E-stained sections. The necrotic core (NC) was defined as the area devoid of nuclei underneath a formed fibrous cap. Collagen content was quantified on Picrosirius Red-stained sections. Vulnerability plaque index (VPI) was calculated as VPI = (% NC area + % CD68 area) / (% SMA area + % collagen area).

**Flow cytometry analysis**

Isolated CCA samples were mixed with digestion buffer, consisting of collagenase type XI (125 U/ml, C7657), hyaluronidase type 1-s (60 U/ml, H3506), DNase I (60 U/ml, D5319), collagenase type I (450 U/ml, C0130; all enzymes from Sigma Aldrich, Germany) in 1x PBS[9], and were digested at 750 rpm for 30 min at 37 °C. After digestion, CCA materials were homogenized through a 40 µm cell strainer, washed at 500 g for 7 min at 4 °C and resuspended in flow cytometry staining buffer (00-4222-26, ThermoFisher) to generate single cell suspensions. Cell suspension were incubated with according flow cytometry antibodies and analysed using a spectral flow cytometer (Northern Light, CYTEK, US). Please find a detailed antibody list for flow cytometry analysis in the resource table below.

**Analysis of plaque-infiltrating leukocytes**

Circulating leukocytes were discriminated by intravenous administration of an anti-CD45 antibody[10] (eFluor450, clone: 30-F11, eBioscience, US) immediately before stroke surgery. 24 hours after stroke, to exclude the blood contamination in the CCA, an anti-CD45 antibody (APC-Cy7, clone: 30-F11, Biolegend, US) was injected intravenously 3 min before sacrifice. Then CD45 eFluor450 positive but APC-Cy7 negative population were considered as the "infiltrating leukocytes" population.

**Immunoblot analysis**

Ipsi- and contralateral CCA materials were carefully isolated and snap frozen on dry ice. Whole frozen CCA was lysed with RIPA lysis/extraction buffer with added protease/phosphatase inhibitor (Thermo Fisher Scientific, US). Total protein quantified using the Pierce BCA protein assay kit (Thermo Fisher Scientific). Whole tissue lysates were fractionated by SDS-PAGE and transferred onto a polyvinylidene difluoride membrane (BioRad, Germany). After blocking for 1 hour in TBS-T (TBS with 0.1 % Tween 20, pH 8.0) containing 4 % skin mile powder (Sigma, Germany), the membrane was washed with TBS-T and incubated with the primary antibodies against following antibodies: mouse anti-caspase-1 (1:1000; AdipoGen, US), rabbit anti-actin (1:1000; Sigma, Germany) and rabbit anti-Factor XII (1:1000, Invitrogen, US). Membranes were washed three times with TBS-T and incubated for 1 hour with HRP-conjugated anti-rabbit or anti-mouse secondary antibodies (1:5000, Dako, Denmark) at room temperature. Membranes were washed three times with TBS-T, developed using ECL substrate (Millipore, US) and acquired via the Vilber Fusion Fx7 imaging system.

**Enzyme Linked Immunosorbent Assay (ELISA)**

CCA tissue samples were carefully isolated and snap frozen on dry ice. Whole frozen CCA was lysed with cell lysis buffer (#895347, R&D system, US). Then, concentration of IL-1β in total CCA lysates was measured by ELISA according to the manufacturer's instructions (MLB00C, R&D system). Absorbance at 450 nm was measured by iMark Microplate reader (BIO-RAD, Germany)

**MMP2 and MMP9 in situ zymography**

MMP2 and MMP9 in situ zymography on CCA sections was performed as previously described with slight modifications[11]. DQ-gelatin (D12054, Invitrogen) was dissolved in reaction buffer ((50 mM Tris-HCl, 150 mM NaCl, 5 mM CaCl2, 200 mM sodium azide, pH 7.6). 5 µm cryosections were incubated for 2 h at 37 °C with the gelatin-containing reaction buffer. Negative control sections were pre-incubated for 1 h with the MMP-inhibitor

1,10-Phenatheroline (Sigma). Nuclei were stained with DAPI. MMP activity was detected with an Axio Observer Z1 microscope with 20 x magnification (Carl Zeiss, Germany). Data is shown as normalized MMP intensity (Normalized MMP area = MMP+ area / intima area).

**Neutrophil isolation and stimulation**

Neutrophils were generated from tibia and femur of transcardially perfused wild type mice. After isolation and dissection of tibia and femur, bone marrow was flushed out of the bones through a 40 µm strainer using a plunger and 1 ml syringe filled with sterile 1 x PBS. Strained bone marrows cells were washed with PBS, and resuspended in 1 x sterile PBS with 5 % BSA. Afterwards, neutrophils were isolated by using neutrophil isolation kit (130-097-658, Miltenyi Biotec) according to the manufacturer's instructions. 1 x $10^7$ cells were plated onto 150 mm culture dishes in RIPA 1640 (Gibco, US), supplemented with 10 % FBS and 1 % penicillin/streptomycin. Cells were stimulated overnight with 100 nM phorbol 12-myristate 13-acetate (PMA) at 37 °C with 5 % $CO_2$.

**Free nucleic acid quantification**

Total circulating DNA, single strand (ss) DNA and double strand (ds) DNA) levels in the plasma of mice and human patients were first purified with Plasma/serum Cell-free circulating DNA purification mini kit (#55100, NORGEN Biotek, Canada), according to the manufacturer's instructions. Afterwards, total DNA and ssDNA were measured with Nanodrop Spectrophotometer (1000ND, Peqlab, US). dsDNA was measured with a Qubit 2.0 fluorophotometer (Invitrogen, US) using a specific fluorescent dye binding dsDNA (HS dsDNA Assay kit, Thermo Fisher Scientific). Dilutions and standards were generated following the manufacturer's instructions.

**Bone Marrow-Derived Macrophages (BMDM) isolation and cell culture**

BMDMs were isolated and cultured as previously described[2]. Briefly, BMDMs were generated from tibia and femur of transcardially perfused wild type mice. After careful isolation and dissection of tibia and femur, bone marrow was flushed out of the bones through a 40 µm strainer using a plunger and 1 ml syringe filled with sterile 1x PBS. Strained bone marrow cells were washed with PBS, and resuspended in DMEM + GlutaMAX-1 (Gibco, US), supplemented with 10 % fetal bovine serum (FBS) and 1 % Gentamycin (Thermo Scientific, US) and counted. 5 x $10^7$ cells were plated onto 150 mm culture dishes. Cells were differentiated into BMDMs over the course of 8-10 days. For the first days after isolation, cells were supplemented with 20 % L929 cell-conditioned medium (LCM), as a source of M-CSF. Cultures were then maintained at 37 °C with 5% $CO_2$ until they reached ≥ 90% confluence.

**BMDM stimulation with sham or stroke serum**

BMDMs were cultured for 8-10 d for full differentiation. Cells were then harvested, washed, counted, and seeded in flat-bottom tissue culture-treated 24-well plates at a density of 2 x $10^5$ cells per well in a total volume of 500 µl, and then cultured overnight for at least 16 h. BMDMs were then stimulated with LPS (100ng/ml) for 4 h. Afterwards the cells were incubated with serum from either stroke or sham operated wild type mice (4h post-surgery) at a concentration of 25 % total volume for 1 h. Control treated BMDMs received only FBS-containing culture medium. After stimulation, the supernatant was discarded and the cells were washed with sterile PBS to ensure no leftover serum on the cells. Afterwards, 500 µl serum-free DMEM was added to the BMDMs, which were then incubated overnight for 16 h at 37 °C and 5 % $CO_2$. The culture medium was then collected for analysis.

**Gelatin zymography of mouse CCA extracts, BMDMs culture medium and patient plaque lysates**

CCA tissue extracts were analyzed using gelatin zymography (Novex TM 10 % zymography plus protein; ZY00100BOX, Thermo Scientific, US) according to the manufacturer's instructions. CCA tissue was lysed with cell lysis buffer (#895347, R&D system, US). Total protein was quantified using Pierce BCA protein assay kit (Thermo Fisher Scientific, US). Aliquots of appropriately diluted tissue extracts were loaded on gels at total volume of 20 µl. After electrophoresis, gels were incubated in 1 x Zymogram Renaturing Buffer (LC2670, Invitrogen, US) for 30 min at room temperature with gentle agitation. Afterwards Zymogram Renaturing Buffer was decanted and 1 x Zymogram Developing Buffer (LC2671, Invitrogen) was added to the gel. The gel was then equilibrated for 30 minutes at room temperature with gentle agitation. After an additional wash with 1x Zymogram developing buffer, the gels were incubated at 37 °C overnight. Gels were stained with colloidal blue staining kit (LC6025, Invitrogen, US) and acquired on a gel scanner. BMDM culture medium was collected and loaded on gelatin zymography gels at total volume of 25µl. MMP activity in BMDM culture medium was analysed using the same protocol as for the tissue samples.

***En face*** **immunofluorescence staining**

Both, ipsi- and contralateral CCA were carefully dissected and adventitial fat and ligation nodes were thoroughly trimmed away. CCAs were then cut open, unfolded, and pinned out on a silicon-elastomer for fixation in 4 % PFA at room temperature for 2 h. The CCAs then were washed for 1h at room temperature in 5 % BSA with 0.3 % Triton-100X (Sigma Aldrich, Germany). Afterwards, CCAs were incubated with rabbit anti-Factor XII (1:100, PA5-116703, Invitrogen, US) at 4 °C overnight. After washing in 5 % BSA with 0.3 % Triton-100X for 1 h at room temperature, CCAs were incubated with AF647 goat anti-rabbit (1:100, Invitrogen, US) and DAPI for 2 h at room temperature. Finally, CCAs were mounted with fluoromount medium (Sigma Aldrich, Germany). Microphotographs were taken with a confocal microscope (LSM 980; C. Zeiss, Germany).

**Statistical analysis**

Data were analysed using GraphPad Prism version 6.0. All summary data are expressed as the mean ± standard deviation (s.d.) unless indicated otherwise. All datasets were tested for normality using the Shapiro-Wilk normality test. The groups containing normally distributed independent data were analysed using a two-way Student's t test (= 2 groups) or ANOVA (for > 2 groups). Normally distributed dependent data were analysed using a 2-way ANOVA. The remaining data were analysed using the Mann-Whitney U test (= 2 groups) or Kruskal-Wallis Test (H test, for > 2 groups). P value were adjusted for comparison of multiple comparisons using Bonferroni correction or Dunn's multiple comparison tests. P values < 0.05 was considered to be statistically significant.

**Resource list**

| REAGENT or RESOURCE | SOURCE | IDENTIFIER |
|---|---|---|
| **Antibodies** | | |
| Anti-CD68 antibody [FA-11], Rat | Abcam | ab53444 |
| Anti-alpha smooth muscle Actin antibody[1A4], Mouse | Abcam | ab7817 |
| Anti Iba-1, Rabbit | Wako | 019-19741 |
| Anti-Ki-67 (D3B5) mAb, Rabbit | Cell Signaling | 9129S |

| | | |
|---|---|---|
| Anti-mouse caspase-1 (p20; CASPER1; mouse) | Adipogen | AG-20B-0042-C100 |
| Recombinant Anti-MMP2 antibody | Abcam | ab92536 |
| Anti-MMP9 antibody | Abcam | ab38898 |
| Anti-mouse actin, Rabbit | Sigma | A2066-.2ml |
| Anti-human caspase-1 (p20, BALLY-1; mouse) | Adipogen | AG-20B-004 |
| Anti-Factor XII Polyclonal Antibody, Rabbit | Invitrogen | PA5-116703 |
| Anti-CD31 Antibody, Rat | OriGene | BM4086 |
| Anti-Von Willebrand Factor Antibody, Sheep | Abcam | ab11713 |
| Anti-CD41 [MWReg30] Antibody, Rat | Abcam | ab33661 |
| Anti-Fibrinogen Antibody, Rabbit | Abcam | ab34269 |
| Goat anti-Rabbit IgG (H+L) Crossed-Adsorbed Secondary Antibody, Alexa Fluor 594 | Invitrogen | A-11005 |
| Goat anti-Mouse IgG (H+L) Highly Cross-Adsorbed Secondary Antibody, Alexa Fluor 488 | Invitrogen | A-32723 |
| Goat anti-Rat IgG (H+L) Cross-Adsorbed Secondary Antibody, Alexa Fluor 647 | Invitrogen | A-21247 |
| Donkey anti-Sheep IgG (H+L) Cross-Adsorbed Secondary Antibody, Alexa Fluor 594 | Invitrogen | A-11016 |
| Anti-mouse IgG (goat, HRP-conjugated) | Dako | P0447 |
| Anti-rabbit IgG (goat, HRP-conjugated) | Dako | PI-1000 |
| Anti-mouse CD45 (APC-Cy7; 30-F11) | Biolegend | 103116 |
| Anti-mouse CD45 (eFluor450; 30-F11) | Invitrogen | 48-0451-82 |
| Anti-mouse CD11b (PerCP-Cy5.5; M/70) | Invitrogen | 45-0112-82 |
| Anti-mouse F4/80 (PE-Cyanie7; BM8) | Invitrogen | 25-4801-82 |
| Anti-mouse Ly6G (PE-Fluor610; 1A8-Ly6g) | Invitrogen | 61-9668-82 |
| Anti-mouse Ly6C (BV570; HK1.4) | Biolegend | 128030 |
| Anti-mouse CD192 (APC; SA203G11) | Biolegend | 150628 |
| Anti-mouse MHC II (PE; NIMR-4) | Invitrogen | 12-5322-81 |
| Anti-human CD3 (FITC; HIT3a) | Invitrogen | 11-0039-42 |
| Anti-human CD8a (PE; SK1) | Invitrogen | 12-0087-42 |
| Anti-human CD19 (APC; HIB19) | Invitrogen | 17-0199-42 |
| Anti-human CD45 (eFluor 450; 2D1) | Invitrogen | 48-9459-42 |
| Anti-human CD14 (PerCP-Cy5.5; 61D3) | Invitrogen | 45-0149-42 |
| Anti-human CD16 (FITC; CB16) | Invitrogen | 11-0168-42 |
| Anti-human CD11b (PE; ICRF44) | Invitrogen | 12-0118-42 |
| **Chemicals, peptides, and recombinant proteins** | | |
| DMEM+GlutaMAX (4.5g/I D-Gluc. / Pyruvate) | Gibco | 31966-021 |
| Fetal calf serum (FCS) | Gibco | 105000-064 |
| Gentamycin (50mg/ml) | Gibco | 15750-045 |

| | | |
|---|---|---|
| RPMI 1640 (L-Glutamine / 25 mM HEPES) | Gibco | 52400-025 |
| Penicilin / Streptomycin | Gibco | 15140-122 |
| LPS from E.coli | Adipogen | IAX-100-013-M001 |
| Isopentane | Sigam | 277258-1L |
| Phorbol 12-myristate 13-acetate (PMA) | Adipogen | AG-CN2-0010-M005 |
| Human recombinant DNase I | Roche | 4716728001 |
| Caspase-1 inhibitor (VX-765, Belnacasan) | Invivogen | Inh-vx765i-1 |
| NLRP3 inflammasome inhibitor (MCC950) | Invivogen | Inh-mcc |
| **REAGENT** | | |
| RIPA lysis buffer (+protease/phosphatase inhibitor) | Thermo Fisher Sci | 8990 |
| Cell lysis buffer 2 | R&D system | 895347 |
| Zymogram Renaturing Buffer | Invitrogen | LC2670 |
| Zymogram Developing Buffer | Invitrogen | LC2671 |
| Collagenase type XI | Sigma | C7657 |
| Hyaluronidase type 1-s | Sigma | H3506 |
| DNase I | Sigma | D5319 |
| Collagenase type I | Sigma | C0130 |
| Ethylenediamine tetraacetic acid (EDTA) | Roth | Art.No.8043.2 |
| Flow cytometry staining buffer | Thermo Fisher Sci | 00-4222-26 |
| Histopaque solution | Sigma | Histopaque-1119 |
| DQ-gelatin | Invitrogen | D12054 |
| **Critical commercial assays** | | |
| Duoset ELISA murine IL-1beta | R&D system | MLB00C |
| Fluoro-Jade C Ready-to-Dilute Staining Kit | Biosensis | TR-100-FJ |
| Click-iT™ Plus TUNEL Assay for In Situ Apoptosis Detection, Alexa Fluor™ 647 dye | Thermo Fisher Sci | C10619 |
| Picro-Sirius Red Stain Kit (Cardiac Muscle) | Abcam | Ab245887 |
| Colloidal blue staining kit | Invitrogen | LC6025 |
| Neutrophil isolation kit | Miltenyi Biotec | 130-097-658 |
| Plasma / Serum Cell-Free Circulating DNA Purification Mini Kit | NORGEN Biotek | 55100 |
| HS dsDNA Assay Kit | Thermo Fisher Sci | Q32851 |
| **Software** | | |
| Cytek Northern lights | Cytek Biosciences | N/A |
| Microsoft Excel | Microsoft Corporation | N/A |
| FlowJo v. 10.6 | Treestar Inc. | N/A |
| GraphPad Prism 6 | Graphpad SoftwareInc | N/A |
